# Crk proteins activate the Rap1 guanine nucleotide exchange factor C3G by segregated adaptor-dependent and -independent mechanisms

**DOI:** 10.1101/2022.11.24.515150

**Authors:** Antonio Rodríguez-Blázquez, Arturo Carabias, Alba Morán-Vaquero, Sergio de Cima, Juan R. Luque-Ortega, Carlos Alfonso, Peter Schuck, José Antonio Manso, Sandra Macedo-Ribeiro, Carmen Guerrero, José M de Pereda

**Affiliations:** Centro de Investigación del Cáncer and Instituto de Biología Molecular y Celular del Cáncer, Consejo Superior de Investigaciones Científicas (CSIC)- Universidad de Salamanca, 37007 Salamanca, Spain; Instituto de Investigación Biomédica de Salamanca (IBSAL), Salamanca, Spain; Centro de Investigaciones Biológicas Margarita Salas, CSIC, Ramiro de Maeztu 9, 28040 Madrid, Spain; Laboratory of Dynamics of Macromolecular Assembly, National Institute of Biomedical Imaging and Bioengineering, National Institutes of Health, Bethesda, Maryland 20892; IBMC-Instituto de Biologia Molecular e Celular, Universidade do Porto, 4200-135 Porto, Portugal; i3S-Instituto de Investigação e Inovação em Saúde, Universidade do Porto, 4200-135 Porto, Portugal; Departamento de Medicina, Universidad de Salamanca, Instituto de Investigación Biomédica de Salamanca (IBSAL), 37007 Salamanca, Spain

**Author notes:** These authors contributed equally to this work (Antonio Rodríguez-Blázquez, Arturo Carabias). Structural Molecular Biology Group, Novo Nordisk Foundation Centre for Protein Research, Faculty of Health and Medical Sciences, University of Copenhagen, Blegdamsvej 3-B, DK- 2200 Copenhagen N, Denmark.

**Keywords:** Ras-associated protein 1, RapGEF1, signal transduction, tyrosine phosphorylation, Src-homology 2 domain, Src-homology 3 domain

## Abstract

C3G is a guanine nucleotide exchange factor (GEF) that activates Rap1 to promote cell adhesion. Resting C3G is autoinhibited and the GEF activity is released by stimuli that signal through tyrosine kinases. Tyrosine phosphorylation of C3G and interaction with Crk adaptor proteins, whose expression is increased in multiple human cancers, participate in C3G activation. However, the molecular details of C3G activation and the interplay between C3G phosphorylation and Crk interaction are poorly understood. Here, we combine biochemical, biophysical, and cell biology approaches to elucidate the mechanisms of C3G activation. CrkL interacts through its SH3N domain with the proline-rich motifs P1 and P2 of inactive C3G in vitro and in Jurkat and HEK293T cells, and these sites are necessary to recruit C3G to the plasma membrane. However, direct stimulation of the GEF activity requires binding of Crk proteins to the P3 and P4 sites. P3 is occluded in resting C3G and is essential for activation, while P4 contributes secondarily towards complete stimulation. Tyrosine phosphorylation of C3G alone causes marginal activation. Instead, phosphorylation primes C3G lowering the concentration of Crk proteins required for activation and increasing the maximum activity. Unexpectedly, optimal activation also requires the interaction of CrkL-SH2 domain with phosphorylated C3G. Phosphorylation and Crk-binding form a two-factor mechanism that ensures tight control of C3G activation. The simultaneous SH2 and SH3N interaction of CrkL with C3G, required for the activation, reveals a novel adaptor-independent function of Crk proteins relevant to understanding their role in physiological signaling and their deregulation in diseases.

## Introduction

C3G (also known as RapGEF1) is a ubiquitously expressed guanine nucleotide exchange factor (GEF) that activates primarily the small GTPases of the Ras family Rap1a and Rap1b [1, 2]. C3G also activates efficiently the Rho family GTPase TC10 [3]. C3G promotes the conformational switch of these GTPases from an inactive state bound to guanosine diphosphate (GDP) to an active state bound to guanosine triphosphate (GTP). Through the stimulation of Rap1 proteins, C3G regulates integrin-mediated cell adhesion and migration [4]. C3G also regulates cadherin-based cell-cell junctions, cell proliferation, differentiation, apoptosis and exocytosis [5, 6].

The sequence of the main variant of human C3G (isoform a, 1077 residues) contains three distinct regions (Fig. 1A). The N-terminal domain (NTD) harbors a binding site for E-cadherin [7]. The central region contains five Pro-rich motifs (PRMs), named P0 to P4, that are binding sites for specific Src homology 3 (SH3) domains; hence this region is known as the SH3-binding domain or SH3b. Finally, the C-terminal catalytic region consists of a Ras exchanger motif (REM) and a Cdc25 homology domain (Cdc25HD) that is responsible for the GEF activity.

**Figure 1.**
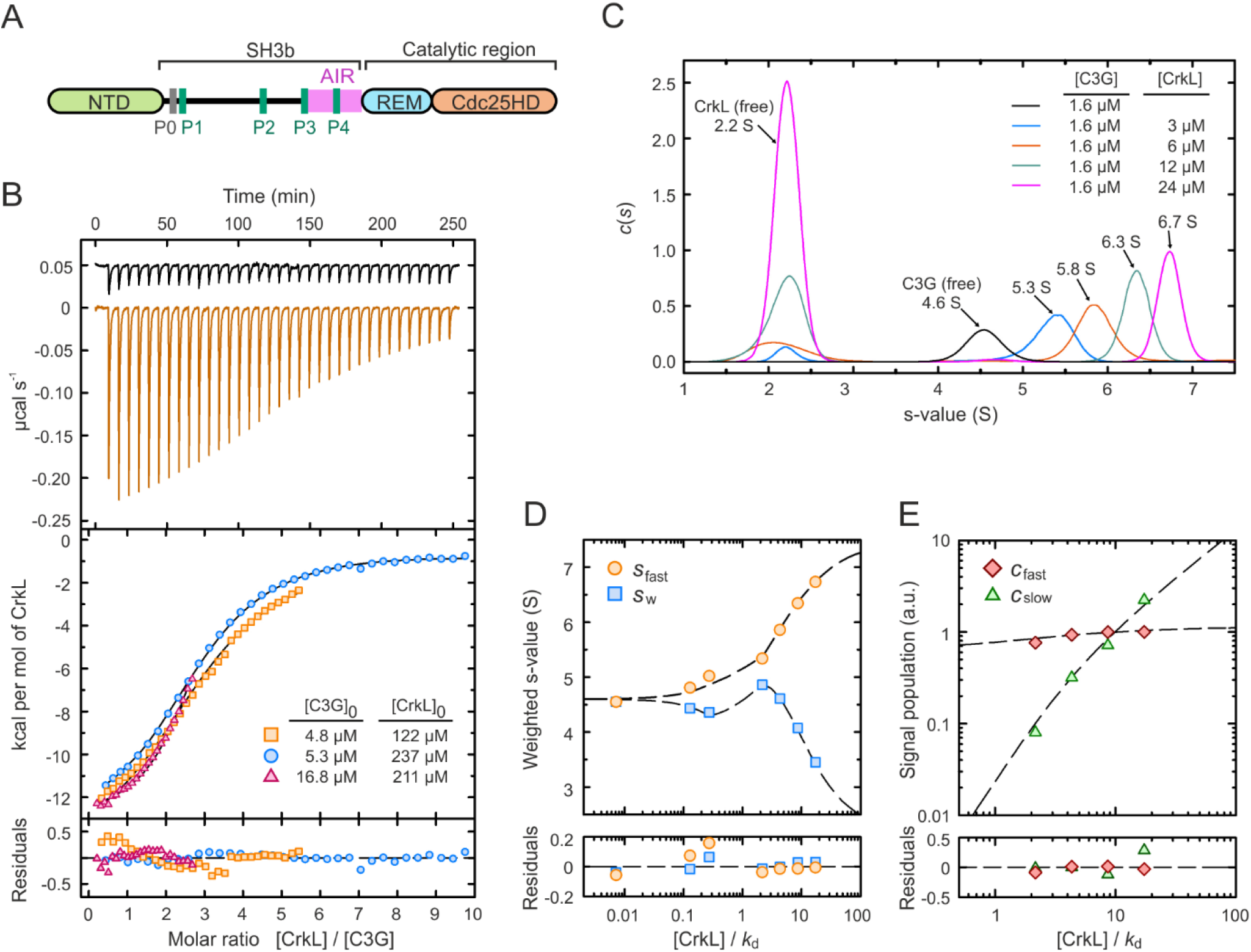
Binding of CrkL to full-length C3G. (**A**) Schematic representation of the domain structure of C3G. (**B**) ITC analysis of CrkL binding to C3G. Upper panel shows thermograms of the dilutions of CrkL in buffer (black line) and of the titration of 4.8 µM C3G with CrkL (orange line). The middle panel shows the binding isotherms of three independent titrations as indicated; lines are the theoretical binding curves obtained by the global fit. Residuals from the fitted model are shown in the lower panel. (**C**) Sedimentation coefficient distributions of C3G alone and in the presence of increasing concentrations of CrkL as indicated. (**D**) Isotherms of the *s*_fast_ obtained from the integration of the fastest *c*(s) peaks in C, and *s*_w_ determined from the integration of the entire *c*(s) distribution. (**E**) Signal amplitudes *c*_fast_ and *c*_slow_ determined by the integration of the *c*(s) peaks. Dashed lines in D and E represent the best fit using the three-sites binding model based on the Gilbert-Jenkins theory.

The activity of C3G is self-regulated by two intramolecular interactions [8]. First, binding of the NTD to the REM domain favors the GEF activity. Second, a segment of the SH3b downstream of the P3 motif constitutes an autoinhibitory region (AIR) that binds to the Cdc25HD and blocks the GEF activity possibly by obstructing the interaction with the GTPase. The initial segment of the AIR (residues 545-569) is sufficient for binding to the catalytic domain but does not block the GEF activity, thus, we refer to it as Cdc25HD-binding region or AIR-CBR. Inhibition of the GEF activity requires a second segment of the AIR (residues 570-646, inhibitory tail or AIR-IT) that by itself is not sufficient to sustain binding to the Cdc25HD. Residues M551, Y554, and M555 in the AIR-CBR are essential for binding to the Cdc25HD. Two of these residues are targeted by somatic missense mutations detected in non-Hodgkin’s lymphomas: Y554H in follicular lymphoma [9] and M555K in diffuse large B cell lymphoma [10]. These mutations disrupt the autoinhibitory AIR/Cdc25HD interaction and cause constitutive activation of C3G-Rap1 in vitro and in cells [8].

Crk adaptor proteins play a major role in the activation of C3G. Mammalian Crk proteins include CrkI and CrkII, which are alternative splicing variants of a single gene, and the related Crk-like (CrkL) protein [11]. CrkII and CrkL have a similar domain organization with an N-terminal Src homology 2 (SH2) domain, and two SH3 domains (SH3N and SH3C), while CrkI only contains the SH2-SH3N domains. Crk SH2 domains bind to phosphorylated tyrosine residues in the context of pY-x-x-P motifs (where pY is phospho-tyrosine) [12]. The SH3N domains bind to the P1, P2, P3, and P4 motifs in the SH3b of C3G, which display a P-x-x-P-X-R/K consensus sequence [13]. Finally, the SH3C domains of Crk proteins are atypical and do not bind to PRMs [14].

C3G is activated by diverse stimuli that include ligation of B and T cell receptors, growth factors, hormones, cytokines and mechanical stretching (reviewed in [5]), which operate through the activation of tyrosine kinases. C3G interacts with CrkII and CrkL in the cytosol of unstimulated cells [15, 16]. Upon stimulation, the Crk/C3G complexes are recruited to signaling sites at the plasma membrane through the binding of the SH2 domain of Crk to phospho-tyrosine-containing proteins such as growth factor receptors, p130Cas, Cbl, and paxillin [17]. C3G is phosphorylated at Y504 and other tyrosine residues during activation [18, 19]. C3G is a substrate for the Src-family kinases Src [20], Hck [21], and Fyn [22], and for c-Abl [19] and its oncogenic version Bcr-Abl [23].

Binding of CrkII or CrkL to C3G directly stimulates the GEF activity in vitro. Similarly, phosphorylation of purified C3G by Src also results in a moderate activation [8, 24]. Binding of Crk proteins and Src-mediated phosphorylation of C3G are independent and additive activating stimuli, both of which are required for the efficient stimulation of the GEF activity [8].

While the general elements of the physiological activation of C3G have been identified, it was still unknown how phosphorylation of C3G and the binding of Crk proteins combine to stimulate the GEF activity of C3G. Here, we present a comprehensive and quantitative characterization of the binding of CrkL to C3G and the activation of C3G and phosphorylated C3G by Crk proteins. We show that tyrosine phosphorylation of C3G mainly contributes to the activation by facilitating and amplifying the Crk-mediated stimulation of the GEF activity. Our results also reveal segregated dual roles of CrkL for the recruitment and activation of C3G. Finally, we show that Crk proteins directly stimulate the GEF activity of C3G through a novel mechanism that is independent of their adaptor role.

## Results

### Binding of CrkL to full-length C3G

To characterize the interaction between CrkL and C3G, first, we analyzed the binding of CrkL to the isolated PRMs using fluorescein-labeled peptides of the P1, P2, P3 and P4 (Fig. S1). CrkL binds to the four PRMs with very similar affinity, the estimated dissociation constants (*k*_d_) ranged between 1.4 and 3.0 µM; which were very similar to the reported affinity of CrkL for the P1 (*k*_d_ 2.8 µM) [25].

Next, we analyzed the binding of CrkL to full-length C3G using isothermal titration calorimetry (ITC). Three independent experiments were done at different starting concentrations of C3G and CrkL to cover a wide range of the binding reaction (Fig. 1B, Table S1). Initially, we analyzed each titration individually with the MicroCal Origin ITC software using an independent-site (non-cooperative) model with no assumptions on the stoichiometry (N), which yielded values of N ∼3 molecules of CrkL per molecule of C3G and *k*_d_ in the range 2.8-3.6 μM. These affinities were similar to those of the binding of CrkL to the isolated PRM peptides (see above). Next, a global simultaneous analysis of the three titrations was performed with the program SEDPHAT. Based on the previous estimation of a 1:3 stoichiometry, we fitted a model with three symmetric binding sites to the experimental isotherms, which yielded a microscopic *k*_d_ of 2.3 μM.

To further characterize the interaction of CrkL with C3G, we used sedimentation velocity (SV) as an orthogonal method. C3G at 1.6 μM sedimented as a discrete peak with a sedimentation coefficient (*s*) of 4.6 S (*s*_20,w_ 4.9 S), which represents 92% of the material (Fig. 1C, Table S2). Complementarily, we used dynamic light scattering (DLS) to determine the diffusion coefficient of C3G (3.36 × 10^−11^ m^2^ s^-1^). The estimated molecular weight of C3G calculated using the sedimentation and diffusion coefficients through the Svedberg equation was 127 kDa, which matched the molecular weight calculated from the sequence (122 kDa). The sedimentation profiles of CrkL at concentrations ranging from 3 to 24 μM displayed major single peaks (>95% monodisperse) with *s* ∼2.2 S (*s*_20,w_ 2.4 S) (Fig. S2). The diffusion coefficient of CrkL determined by DLS was 5.94 × 10^−11^ m^2^ s^-1^ (Table S2). The estimated molecular weight of CrkL obtained from the sedimentation and diffusion coefficients was 34.8 kDa, which coincides with the theoretical mass of the CrkL monomer (33.7 kDa). In summary, C3G and CrkL are monodisperse monomers under these conditions, which allowed the study of their interaction by SV.

Next, we analyzed the sedimentation of mixtures of C3G and CrkL at different concentration ratios (Fig. 1C). Increasing concentrations of CrkL resulted in a progressive displacement of the fast-sedimentation peak from ∼5.3 S to ∼6.7 S. This behavior is typical of rapid interacting systems (dissociation rate constant *k*_off_ > 10^−3^ s^-1^), in which the peak that increased its s-value corresponds to the reaction boundary formed by the CrkL/C3G complex and the free components immediately after dissociating and before associating to form the complex again [26]. Isotherms built from integration of the sedimentation coefficient of the fastest species (*s*_fast_), the weight-average sedimentation coefficient (*s*_w_) (Fig. 1D), and the signal amplitudes of the fast and slow *c*(s) peaks (Gilbert partial concentrations isotherm) (Fig. 1E) were used to fit a model of three symmetric binding sites, yielding a microscopic *k*_d_ value of 1.4 μM, which was similar to the value estimated by ITC (Table S3). We also performed a global and simultaneous analysis of the isotherms extracted from ITC and SV. Using the aforementioned model of three symmetric binding sites, the highest goodness-of-fit predicted a microscopic *k*_d_ value of 2.3 μM (Table S3).

Collectively, ITC and SV data support the notion that on average three molecules of CrkL bind to C3G with low micromolar affinity and that the interaction is highly dynamic.

### P3 is a cryptic CrkL-binding site in unstimulated C3G that is exposed during activation

The previous results suggested that some of the four CrkL-binding sites might not be accessible in resting C3G, which is autoinhibited. To address this question, we analyzed the binding of CrkL to each of the four individual PRMs in full-length C3G. The CrkL-binding sites were disrupted by mutating in each PRM five residues that are essential for the binding to the SH3N domain of Crk [25, 27] (Fig. 2A). We created four mutants, each containing a single unmodified PRM and carrying mutations in the other three. These C3G mutants were named PAAA, APAA, AAPA, and AAAP, to depict whether the P1, P2, P3, or P4 maintained the wild type sequence (P) or carried the mutations to Ala (A). The interaction of these mutants with GST-CrkL-SH3N was analyzed by pull-down assays (Fig. 2B). C3G mutants displaying only the P1 (C3G-PAAA), P2 (C3G-APAA), or P4 (AAAP) bound to CrkL-SH3N, albeit C3G-AAAP was pulled down slightly less than the mutants PAAA and APAA. The interaction of these single-PRM mutants with GST-CrkL-SH3N was lower than that of wild type C3G, suggesting that multivalent contacts with immobilized GST-CrkL-SH3N strengthen the interaction. Finally, almost no interaction was observed with C3G-AAPA.

**Figure 2.**
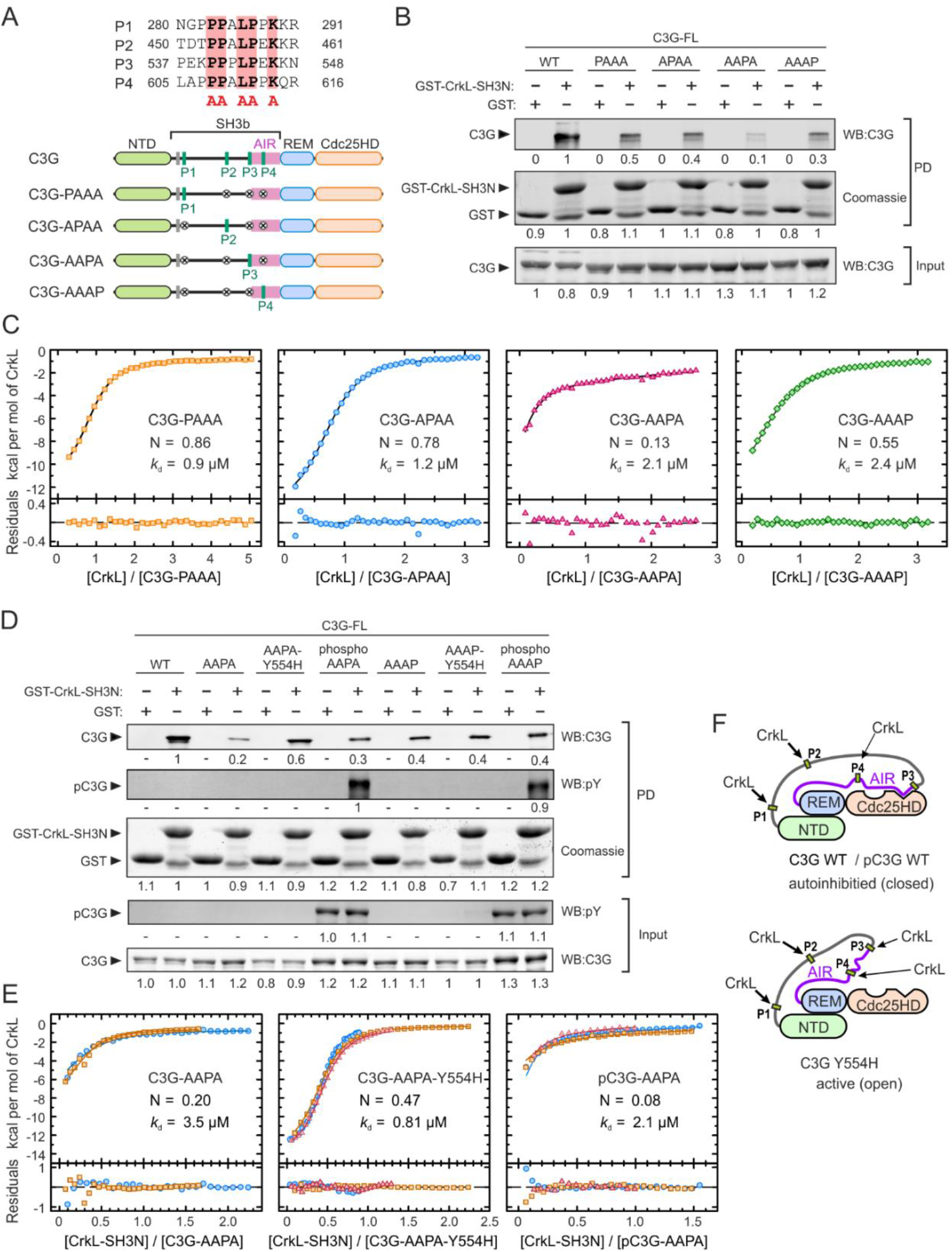
Binding of CrkL to individual sites in full-length C3G. (**A**) Sequences of the four PRMs of human C3G that are binding sites for CrkL. The five key residues in each motif that were mutated to Ala to disrupt CrkL binding are marked by red boxes. Schematic representation of the C3G mutants, each containing a single wild type PRM. (**B**) Analysis by pull-down (PD) of the interaction of GST-CrkL-SH3N with C3G full-length (FL) wild type (WT) and mutants. C3G was detected in the PD and input samples by Western blot (WB). GST-CrkL-SH3N and GST (used as control) were visualized by coomassie staining. Numbers are the relative quantification within each panel. (**C**) ITC analysis of the binding of full-length CrkL to single-PRM mutants of C3G. The binding isotherms (1:1 binding models) and the residuals from the fitted model are shown. N is the estimated competent fraction that reflects the fraction of sites accessible for binding. The corresponding thermograms are shown in Fig. S3A. (**D**) PD analysis of the binding of GST-CrkL-SH3N to C3G WT and single-PRM mutants, in combination with the activating mutation Y554H or after phosphorylation by Src. Numbers are relative quantification of bands as in B. (**E**) ITC binding isotherms of the interaction of the SH3N domain of CrkL to C3G-AAPA, C3G-AAPA-Y554H, and Src-phosphorylated pC3G-AAPA. Thermograms are shown in Fig. S3B. (**F**) Schematic models of the structure of autoinhibited C3G WT and the constitutive active mutant C3G Y554H. The P3 is exposed in the active conformation.

To better understand the differences in binding of CrkL to each PRM in full-length C3G we analyzed the interactions by ITC (Fig. 2C and Table S4). The single-site C3G mutants were titrated with full-length CrkL and the data were analyzed using a 1:1 hetero-association model. CrkL bound to the P1 (C3G-PAAA) and P2 (C3G-APAA) motifs with estimated *k*_d_ ∼1 μM and competent fractions of C3G between 0.8-0.9. The thermograms of CrkL binding to the P3 (C3G-AAPA) and P4 (C3G-AAAP) showed smaller heat signals than the binding to the PAAA and APAA mutants (Fig. S3A). The affinities of CrkL for the P3 and P4 (*k*_d_ ∼2 μM) were only slightly lower than for the P1 and P2. Yet, the estimated competent fractions of C3G-AAPA and C3G-AAAP were 0.13 and 0.55, respectively.

The P3 is juxtaposed to the AIR, and the P4 is within the AIR (Fig. 2A). Therefore, the accessibility to these sites might be linked to the AIR/Cdc25HD autoinhibitory interaction, and accordingly, to the activation state of C3G. To test this hypothesis, we analyzed the binding of CrkL to the P3 and P4 in the presence of the Y554H substitution that disrupts the AIR/Cdc25HD interaction and causes constitutive activation of C3G [8]. The active mutant C3G-AAPA-Y554H was pulled down by GST-CrkL-SH3N more efficiently than the autoinhibited C3G-AAPA (Fig. 2D), while introduction of Y554H in C3G-AAAP did not increase binding to GST-CrkL-SH3N. We also analyzed the effect of the phosphorylation of these C3G mutants with Src, which causes a partial activation of C3G but does not disrupt the AIR/Cdc25HD interaction [8]. Phosphorylated C3G-AAPA was pulled down slightly more than the unphosphorylated protein, although much less than C3G-AAPA-Y554H. In contrast, phosphorylation of C3G-AAAP did not increase the interaction with GST-SH3N.

Next, we used ITC to analyze quantitatively the effect of tyrosine phosphorylation and the Y554H mutation on the binding of CrkL-SH3N to the P3 and P4 motifs (Fig. 2E, Fig. S3B, C and Table S4). The SH3N domain of CrkL bound to the P3 (C3G-AAPA) with comparable affinity (*k*_d_ 3.5 μM) and stoichiometry (N 0.2) as observed for the binding of full-length CrkL (see above). Introduction of the activating mutation Y554H increased slightly the affinity of CrkL-SH3N for binding to the P3 (*k*_d_ 0.8 μM). The main effect of Y554H was the increase of the competent fraction of binding sites to ∼0.5 molecules of SH3N per molecule of C3G-AAPA-Y554H. In contrast, phosphorylation of C3G-AAPA with Src did not increase the fraction of active binding sites (N ∼0.1) or the affinity of CrkL-SH3N with respect to the unphosphorylated protein.

CrkL-SH3N bound to the P4 site in the unphosphorylated and phosphorylated forms of the C3G-AAAP mutant and in the constitutively active form C3G-AAAP-Y554H with minor differences (*k*_d_ 2.4-2.7 μM and N 0.6-0.7) (Fig. S3C), which were also similar to the binding of full-length CrkL to C3G-AAAP (Fig. 2C).

Collectively, the results suggest that in autoinhibited C3G the P1 and P2 sites are constitutively accessible to CrkL. The P4 site is partially exposed in the autoinhibited, active, and Src-phosphorylated states of C3G. Finally, access of the SH3N domain of CrkL to the P3 site depends on the activation state of C3G; P3 is mostly occluded in autoinhibited and phosphorylated (partially active) C3G, and only becomes exposed upon disruption of the AIR/Cdc25HD autoinhibitory interaction (Fig. 2F).

### Binding of CrkL to the P3 and P4 sites is sufficient and necessary for stimulation of C3G GEF activity

The different interaction of CrkL with the PRMs prompted us to study the role of CrkL binding to each PRM in the activation of C3G. Using independently produced proteins of the AIR and Cdc25HD, we had shown that binding of CrkL to the P3 displaces the interaction of the AIR with the Cdc25HD [8]. Here, we characterized the activation of full-length C3G by CrkL. We analyzed mutants containing only one wild type PRM or various combinations of unmodified PRMs (Fig. 3A); we measured their GEF activity using an in vitro fluorescence assay based on the release of the fluorescent GDP analogue mant-dGDP bound to Rap1b (Fig. 3B). The apparent nucleotide dissociation rate constant (*k*_obs_), which is a measure of the GEF activity, was determined for C3G wild type and mutants, alone and in the presence of an excess of CrkL. All mutants showed basal activities similar to that of autoinhibited wild type C3G (Fig. 3C); supporting the notion that the PRMs are not necessary for autoinhibition. CrkL did not activate C3G-PPAA above the basal level, but activated C3G-AAPP to a similar level as wild type C3G. CrkL also stimulated the activity of the mutants C3G-AAPA and C3G-PPPA, but to a level significantly lower than wild type C3G and the AAPP mutant. CrkL did not activate C3G-AAAP; similarly, mutation of the P3 alone, in C3G-PPAP, was sufficient to prevent activation by CrkL.

**Figure 3.**
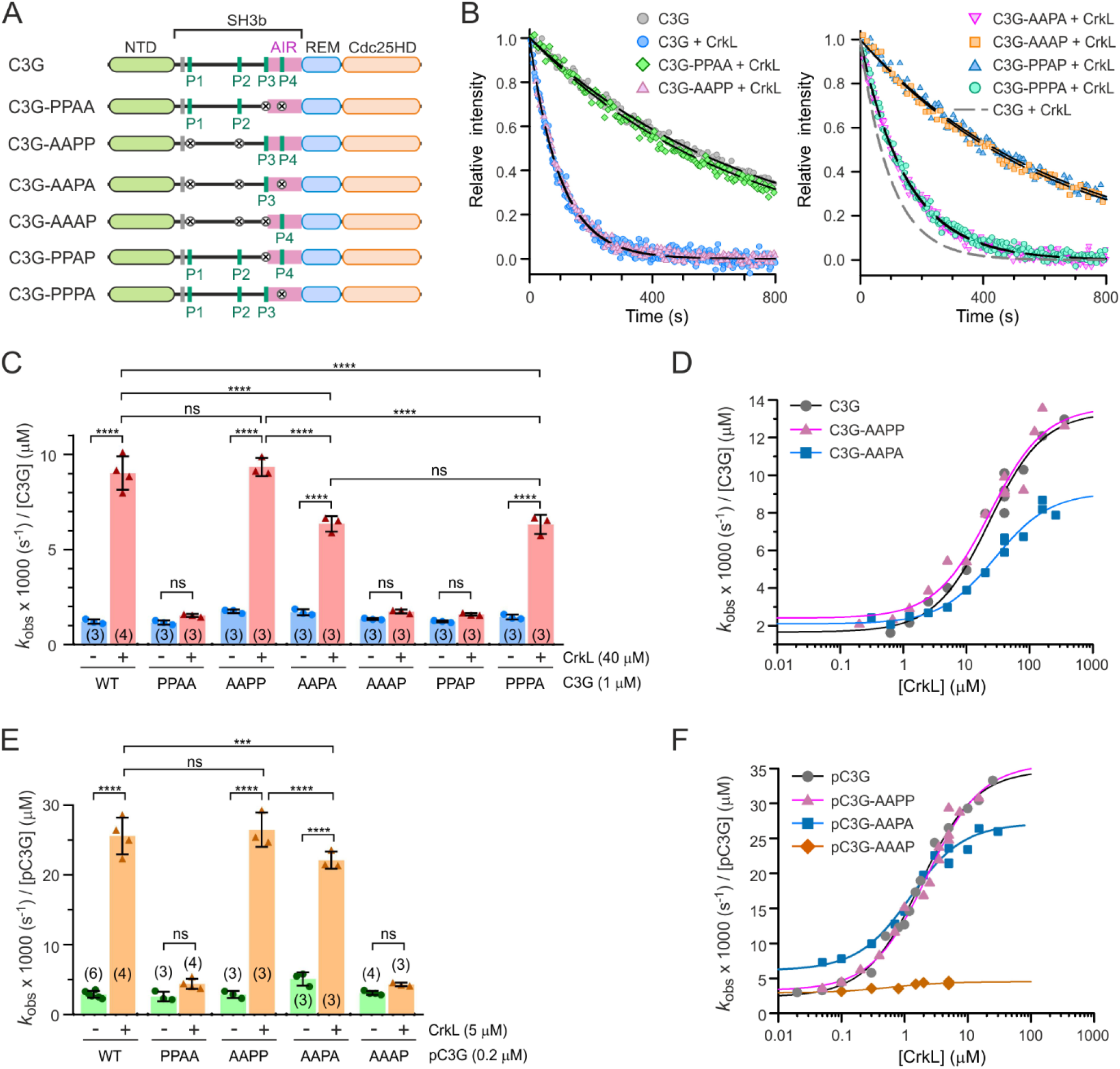
Contribution of individual PRMs to the activation of C3G by CrkL. (**A**) Schematic representation of the PRM mutants used to study the activation of C3G by CrkL. (**B**) Representative exchange reactions of Rap1:mant-dGDP (200 nM) catalyzed by C3G wild type and mutants (1 µM) in the presence of CrkL (40 µM). Lines are the single exponential decay models fitted to obtain the *k*_obs_. (**C**) Nucleotide exchange rates of C3G wild type and mutants (1 µM) alone and in the presence of 40 µM CrkL. Data are shown as scatter plots with bars, means ± standard deviation. The number of independent measurements is indicated in parentheses. Statistical comparison was analyzed using ANOVA followed by Tukey’s multiple comparisons test; *** *P* < 0.001, **** *P* < 0.0001, ns *P* > 0.05. (**D**) Dose-dependent effect of CrkL on the GEF activity of C3G (1 µM) wild type and two mutants. Lines are the fitted sigmoidal models. (**E**) Nucleotide exchange rates of Src-phosphorylated C3G (pC3G, 0.2 µM) wild type and mutants, alone and in the presence of 5 µM CrkL. Statistical comparisons were done as in C. (**F**) Dose-dependent effect of CrkL on the GEF activity of pC3G (0.2 µM) wild type and mutants. Dissociation rate constants in C-F are referred to 1 µM C3G for comparison.

Next, we analyzed the dependence of the activity of C3G wild type and the mutants AAPP and AAPA, with the concentration of CrkL (Fig. 3D). From these dose-response experiments the parameters *k*_free_, *k*_max_, and AC_50_ were estimated. The *k*_max_ is a measure of the maximum activity that CrkL can induce, while the AC_50_ indicates at which concentration of CrkL occurs the midpoint activation. CrkL activated similarly C3G wild type and C3G-AAPP, with AC_50_ ∼23 µM and *k*_max_ ∼13.5 × 10^−3^ s^-1^ μM^-1^, the latter corresponds to an increase between 5.7 and 7.9 of the basal activity, as measured by the *k*_max_/*k*_free_ ratio (Table 1). CrkL also activated C3G-AAPA with a similar AC_50_ of ∼28 µM, yet it induced a lower *k*_max_ of ∼9 × 10^−3^ s^-1^ μM^-1^ that corresponds to a 4.3-fold activation. Collectively, the data revealed that the P1 and P2 do not participate in the direct activation of C3G by CrkL; binding of CrkL to the P3 is sufficient and necessary for activation, but efficient stimulation also requires the interaction of CrkL with the P4.

**Table 1.** Parameters of dose-response activation of C3G by Crk proteins.

| C3G | Crk protein | $\text{AC}_{50} (\mu\text{M})^a$ | $k_{\text{free}} \times 1000 (\text{s}^{-1} \mu\text{M}^{-1})$ | $k_{\text{max}} \times 1000 (\text{s}^{-1} \mu\text{M}^{-1})$ | $k_{\text{max}}/k_{\text{free}}$ |
| --- | --- | --- | --- | --- | --- |
| C3G | CrkL | $22.4 \pm 4.5$ | $1.7 \pm 0.4$ | $13.4 \pm 0.6$ | $7.9 \pm 2.1$ |
| C3G-AAPP | CrkL | $23.7 \pm 6.6$ | $2.4 \pm 0.5$ | $13.6 \pm 0.8$ | $5.7 \pm 1.2$ |
| C3G-AAPA | CrkL | $28.1 \pm 5.7$ | $2.1 \pm 0.2$ | $9.1 \pm 0.4$ | $4.3 \pm 0.5$ |
| C3G | CrkL-SH3N | $26.8 \pm 6.5$ | $1.5 \pm 0.1$ | $6.3 \pm 0.4$ | $4.2 \pm 0.4$ |
| C3G | CrkL-SH2-SH3N | $15.8 \pm 2.6$ | $1.2 \pm 0.2$ | $8.4 \pm 0.2$ | $7.0 \pm 1.3$ |
| C3G | CrkL-SH3N-SH3C | $19.8 \pm 3.9$ | $1.6 \pm 0.3$ | $12.9 \pm 0.6$ | $8.1 \pm 1.7$ |
| C3G | CrkII | $65.7 \pm 20.4$ | $1.6 \pm 0.2$ | $7.2 \pm 0.5$ | $4.5 \pm 0.7$ |
| C3G | CrkL-II-L | $16.7 \pm 4.5$ | $0.9 \pm 0.3$ | $7.9 \pm 0.3$ | $8.8 \pm 3.4$ |
| C3G | CrkII-L-II | $18.7 \pm 2.4$ | $1.2 \pm 0.1$ | $7.9 \pm 0.2$ | $6.6 \pm 0.8$ |
| pC3G | CrkL | $1.7 \pm 0.2$ | $2.3 \pm 0.6$ | $34.2 \pm 1.0$ | $14.9 \pm 4.0$ |
| pC3G-AAPP | CrkL | $2.2 \pm 0.5$ | $3.3 \pm 1.0$ | $35.7 \pm 2.0$ | $10.8 \pm 3.3$ |
| pC3G-AAPA | CrkL | $1.3 \pm 0.3$ | $6.2 \pm 0.9$ | $26.9 \pm 0.8$ | $4.3 \pm 0.6$ |
| pC3G-AAAP | CrkL | $0.6 \pm 0.6$ | $2.9 \pm 0.4$ | $4.5 \pm 0.2$ | $1.6 \pm 0.2$ |
| pC3G | CrkL-SH3N | $12.8 \pm 4.6$ | $2.9 \pm 0.5$ | $13.9 \pm 1.2$ | $4.8 \pm 0.9$ |
| pC3G | CrkL-SH2-SH3N | $4.9 \pm 0.8$ | $2.5 \pm 0.8$ | $37.7 \pm 1.5$ | $15.1 \pm 4.6$ |
| pC3G | CrkL-SH3N-SH3C | $4.5 \pm 1.0$ | $3.7 \pm 0.4$ | $13.4 \pm 0.4$ | $3.6 \pm 0.4$ |
| pC3G | CrkII | $2.6 \pm 0.6$ | $2.1 \pm 0.8$ | $22.3 \pm 0.9$ | $10.6 \pm 4.2$ |
| pC3G | CrkL-II-L | $2.3 \pm 0.5$ | $2.2 \pm 1.3$ | $33.6 \pm 1.2$ | $15.3 \pm 8.5$ |
| pC3G | CrkII-L-II | $1.6 \pm 0.4$ | $2.4 \pm 1.6$ | $30.0 \pm 0.9$ | $12.5 \pm 8.5$ |

<sup>a</sup> Parameters are presented as the fitted value ± the associated standard errors of the nonlinear regression fit.

C3G is phosphorylated in tyrosine residues during activation [18], and tyrosine phosphorylation of C3G and binding of CrkL are additive stimuli that directly activate C3G [8]. Therefore, we also analyzed the activation of phosphorylated-C3G (pC3G) when CrkL binds to different PRMs. C3G wild type and mutants were phosphorylated with Src and their GEF activity was measured alone and in the presence of an excess of CrkL (Fig. 3E). Experiments were done at 0.2 μM pC3G to work below the AC_50_ of CrkL (see below). Data are presented as the specific activity (*k*_obs_) per 1 µM pC3G for comparison with the activity of unphosphorylated proteins. In this regard, the activities of pC3G alone and in the presence of an excess of CrkL were directly proportional to the concentration of C3G in this range of protein concentration (Fig. S4), as is the activity of unphosphorylated C3G [8]. Phosphorylation induced a moderate ∼2-fold increase in the GEF activity of C3G and 5 µM CrkL caused an additional ∼9-fold activation, in agreement with our previous observations [8]. CrkL stimulated pC3G-AAPP to a similar extent as wild type pC3G. CrkL also stimulated pC3G-AAPA, yet the activity induced by CrkL was slightly but significantly lower than that observed for pC3G wild type and pC3G-AAPP. Finally, CrkL did not activate the pC3G-PPAA and pC3G-AAAP mutants.

We also analyzed the dependence of the GEF activity of pC3G, wild type and PRM mutants, with the concentration of CrkL (Fig. 3F). Wild type pC3G was stimulated at lower concentrations of CrkL than the unphosphorylated protein, as shown by a ∼10-fold reduction of the AC_50_ upon phosphorylation (Table 1). In addition, CrkL induced a maximal activity of pC3G that was ∼2.6 times higher than that induced on unphosphorylated C3G. Activation of pC3G-AAPP by CrkL was as that of wild type pC3G. CrkL also activated pC3G-AAPA with a similar AC_50_; yet the maximum activity induced by CrkL was ∼25% lower than for the wild type and the AAPP mutant. Finally, pC3G-AAAP was not stimulated by CrkL.

Collectively, our data revealed that only the P3 and P4 motifs are necessary and sufficient for activation of pC3G by CrkL, with P3 being the main activation site. Phosphorylation of C3G by Src increases both the sensitivity to CrkL-mediated direct activation and the maximal exchange activity induced by CrkL.

### The SH2 and SH3C domains of CrkL also contribute to the activation of C3G

Because CrkL binds to the PRMs P1 to P4 of C3G through the SH3N domain [13], we explored if the CrkL-SH3N domain is sufficient to activate C3G. To that end, the GEF activity of C3G was titrated with several constructs of CrkL (Fig. 4A,B, Table 1). The isolated SH3N domain stimulated C3G to a maximum activity (*k*_max_ 6.3 × 10^−3^ s^-1^ μM^-1^) that was half of that induced by full-length CrkL. This prompted us to analyze the contribution of the SH2 and SH3C domains to the activation of C3G. The maximum GEF activity induced by the fragment SH2-SH3N (*k*_max_ 8.4 × 10^−3^ s^-1^ μM^-1^) was slightly higher than that of the SH3N domain, yet it was lower than the activation observed with full-length CrkL. A fragment encompassing the two SH3 domains (SH3N-SH3C) activated C3G as the full-length protein. Next, we analyzed if the differences in the activation of C3G by the CrkL fragments could be related to their affinity for C3G. We analyzed by ITC the binding of the constructs SH3N, SH2-SH3N, and SH3N-SH3C to full-length C3G (Fig. S5A-C, Table S5). The three fragments of CrkL bound to C3G with similar affinity (*k*_d_ 1.1 - 2.2 µM) as full-length CrkL (see above). The similar affinities of CrkL and its fragments for C3G correlate with comparable AC_50_ values (16-27 µM) for the activation of the GEF activity. In summary, the data suggest that binding of the SH3N to C3G is not modulated by other domains of CrkL; yet, regions outside the SH3N, mainly the SH3C and the linker connecting the two SH3 domains, contribute to the efficacy of the activation of C3G.

**Figure 4.**
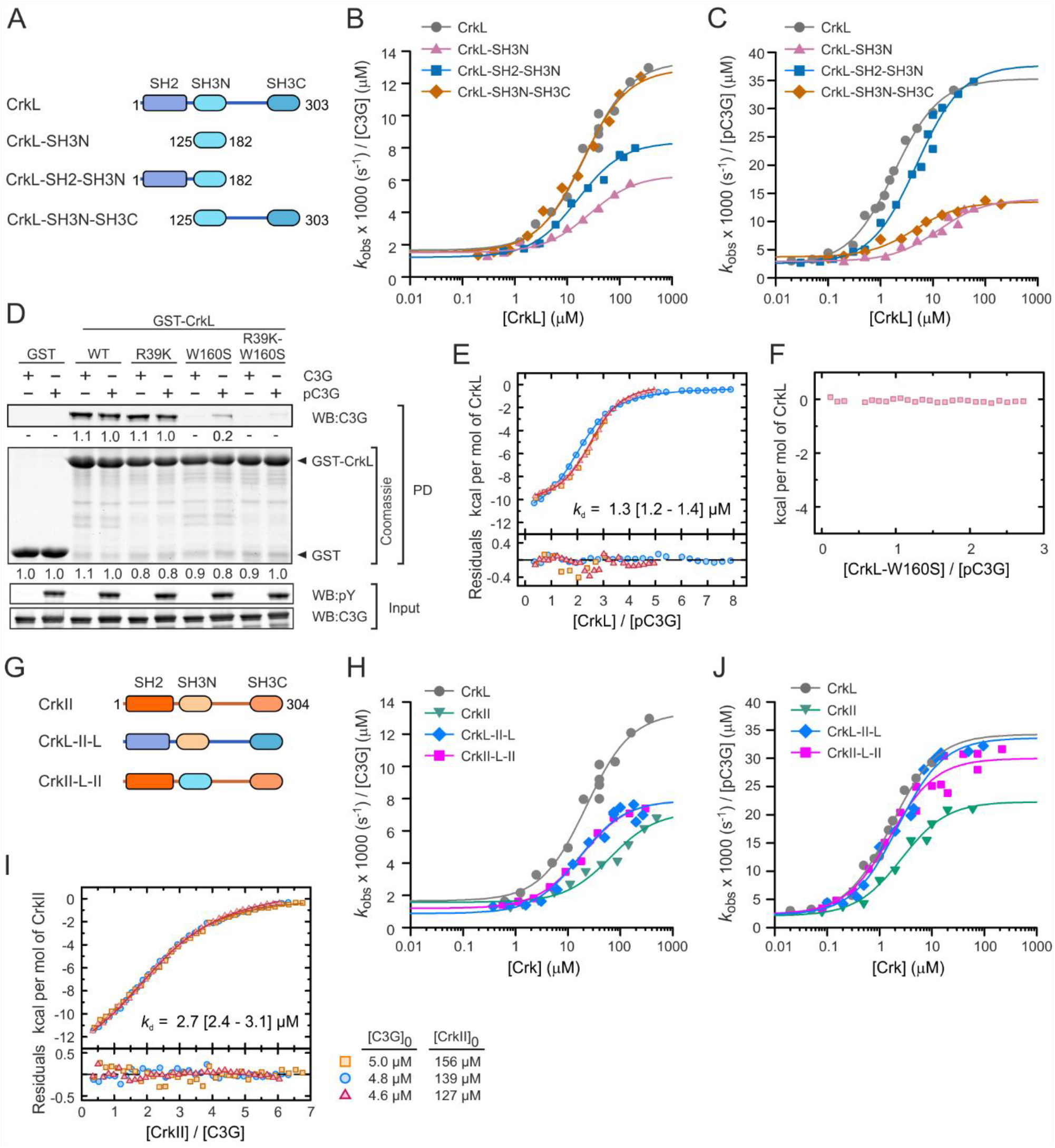
Activation of C3G by CrkL, CrkII, and their constituent domains. (**A**) Schematic representation of the domain structure of CrkL and the deletion mutants analyzed. (**B**) Analysis of the dose-dependent activation of C3G (1 µM) by CrkL full-length and the indicated fragments. Lines in B, C, H, and J are the fitted sigmoidal activation models. (**C**) Dose-dependent activation of Src-phosphorylated C3G (pC3G, 0.2 µM) by CrkL and its deletion mutants. (**D**) Pull-down (PD) analysis of the binding of GST-CrkL, wild type and mutants, to C3G and pC3G. C3G was detected in the PD and in the input samples by Western blot (WB). Phosphorylated C3G was detected with an antibody that recognizes phospho-Tyr (pY). Numbers are the relative quantitation of the bands. (**E**) ITC isotherms of the binding of CrkL to pC3G, three independent titrations are shown. The thermogram of a representative experiment is shown in Fig. S5D. (**F**) ITC analysis of the titration of pC3G with the SH3N-inactive mutant W160S of CrkL. No signal was detected. The corresponding thermogram is shown in Fig. S5E. (**G**) Schematic representation of the domain structure of CrkII and the CrkL-CrkII chimeric proteins. (**H**) Dose-dependent analysis of the activation of C3G (1 μM) by CrkII and the CrkII-CrkL chimeric proteins. (**I**) ITC isotherms (three independent titrations) of the binding of CrkII to C3G. The thermogram of a representative experiment is shown in Fig. S5F. (**J**) Dose-dependent analysis of the activation of pC3G (0.2 μM) by CrkII and the CrkII-CrkL chimeric proteins. Rate constants in B, C, H and J are shown as the specific activities per 1 μM C3G for comparison.

We also analyzed the role of the SH2 and SH3C domains on the activation of C3G phosphorylated by Src (Fig. 4C, Table 1). On the one hand, the isolated SH3N domain and the fragment SH3N-SH3C activated pC3G to a similar maximum activity (*k*_max_ ∼13 × 10^−3^ s^-1^ µM^-1^) that was ∼40% of the maximum activity induced by full-length CrkL. The SH3N alone showed a higher AC_50_ (∼13 µM) than the fragment SH3N-SH3C (AC_50_ 4.5 µM). On the other hand, the SH2-SH3N region activated pC3G to a similar maximum activity (*k*_max_ 37.7 × 10^−3^ s^-1^ µM^-1^) as wild type CrkL. Yet, the SH2-SH3N did not fully recapitulate the activation of pC3G by CrkL, because the SH2-SH3N showed slightly higher AC_50_ (4.9 µM) than full-length CrkL (AC_50_ 1.7 µM), suggesting that the SH3C domain of CrkL also contributes to the efficient activation of pC3G.

Because the SH2 domain of CrkL binds to phospho-tyrosine sites, we analyzed if this domain could bind to Src-phosphorylated C3G. First, we analyzed the interaction using pull-down assays with CrkL fused to GST (Fig. 4D). Wild type GST-CrkL pulled down C3G and pC3G with similar efficiency. To prevent the interaction between the SH3N domain and the PRMs in C3G, we introduced the point substitution W160S in CrkL, which disrupts the PRM-binding site in the SH3N (Fig. S1C). GST-CrkL-W160S did not interact with unphosphorylated C3G but pulled down a small amount of pC3G.

Introduction in CrkL of a second substitution R39K that disrupts the phospho-Tyr binding site, in combination with W160S, prevented the interaction of CrkL with pC3G, supporting the notion that the SH2 domain binds to pC3G. The mutation R93K alone did not reduced the interaction, compared to the wild type GST-CrkL. Next, we used ITC to further analyze the interaction of CrkL with pC3G by an independent method (Fig. 4E,F, Fig. S5D,E, and Table S5). CrkL bound to pC3G with similar affinity (*k*_d_ 1.3 μM) as observed for the binding to unphosphorylated C3G (see above). When pC3G was titrated with CrkL-W160S, no heat-exchange signal was observed (Fig. 4F and Fig. S5E), suggesting that there is no detectable binding in the conditions assayed and that the interaction observed in the pull-downs is of low affinity.

Collectively, our data showed that, in addition to the SH3N, the SH2 and SH3C domains of CrkL contribute to the stimulation of the GEF activity of C3G, and that the SH2 domain plays a key role in the activation of tyrosine phosphorylated C3G possibly through a low affinity interaction with phospho-sites in pC3G.

### C3G is differentially activated by CrkL and CrkII proteins

CrkII also stimulates directly the GEF activity of C3G in vitro [24]. To characterize the activation of C3G by CrkII, first we titrated the GEF activity of unphosphorylated C3G with increasing concentrations of CrkII (Fig. 4G, H, Table 1). CrkII was a weaker activator of C3G than CrkL. The maximum GEF activity of C3G induced by CrkII (*k*_max_ 7.2 × 10^−3^ s^-1^ μM^-1^) was about half of that induced by CrkL. In addition, the half-maximum activity of C3G was reached at a three-fold higher concentration of CrkII (AC_50_ 66 µM) than CrkL. To explore if the lower activation efficiency of CrkII could be related to differences in the interaction with C3G, we analyzed the binding of CrkII to C3G by ITC (Fig. 4I, Fig. S5F, Table S5). CrkII bound to C3G with similar affinity as CrkL.

Because the SH2 and SH3C domains of CrkL contribute to the activation of C3G, we analyzed if the weaker activation of C3G by CrkII could be related to different contributions of the regions outside the SH3N domain. To that end, we created a chimeric protein in which the SH3N domain of CrkL was replaced by the SH3N domain of CrkII, and an equivalent CrkII chimeric mutant that contains the SH3N domain of CrkL. We named these proteins CrkL-II-L and CrkII-L-II to denote if the first (SH2), second (SH3N), and third (SH3C) domains correspond to CrkII or CrkL (Fig. 4G). Analysis of the dose-response activation of unphosphorylated C3G revealed that the two chimeric proteins stimulated C3G in a very similar manner (Fig. 4H). They showed similar AC_50_ values (∼17 µM) as observed for CrkL. Yet, the chimeric proteins induced maximum activities that were about half of the activity induced by CrkL, and were comparable to the *k*_max_ caused by CrkII and the SH2-SH3 fragment of CrkL.

Next, we analyzed the activation of pC3G by CrkII and the CrkII-CrkL chimeric proteins (Fig. 4J, Table 1). CrkII, CrkL-II-L, and CrkII-L-II stimulated pC3G with similar AC_50_ values (1.6 to 2.6 µM) as CrkL. The two chimeric proteins induced maximum GEF activities (*k*_max_ 30-34 × 10^−3^ s^-1^ µM^-1^) that were comparable to that obtained with CrkL, while CrkII was less efficient and stimulated pC3G to ∼60% of the maximum activity of CrkL (Fig. 4J). In summary, our data suggest that the reduced capacity of CrkII to activate pC3G is not due to differences in its individual SH2, SH3N and SH3C domains, because when combined with domains of CrkL they support efficient activation. Therefore, the different activation of C3G may be caused by differences in the inter-domain arrangements of the two Crk proteins.

### C3G recruitment to the plasma membrane requires the P1 and P2 motifs and it is necessary for C3G activation in Jurkat cells

We found that the PRMs P1 and P2 are readily accessible for CrkL binding to autoinhibited C3G in vitro, and these two sites are dispensable for the direct activation of C3G by CrkL. Therefore, we reasoned that the P1 and P2 might participate in the Crk-mediated recruitment of C3G to the plasma membrane during activation. To analyzed the role of the PRMs in the interaction of C3G with CrkL in a cellular model we used Jurkat cells, where CrkL associates with C3G [28]. We used lentiviral transfection to create Jurkat cells stably expressing C3G, wild type and PRM mutants, fused to monomeric enhanced green fluorescent protein (mEGFP). We analyzed the interaction of CrkL with C3G-mEGFP by co-immunoprecipitation (coIP) using affinity resin against GFP (Fig. 5A). CrkL coIP with wild type C3G-mEGFP and with the mutant containing the P1 and P2 (C3G-PPAA-mEGFP), but only a faint presence of CrkL was detected in IPs of cells expressing C3G-AAPP-mEGFP. CrkL also coIP with the single PRM mutants C3G-PAAA-mEGFP and C3G-APAA-mEGFP, indicating that CrkL engages with both the P1 and P2 motifs in Jurkat cells.

**Figure 5.**
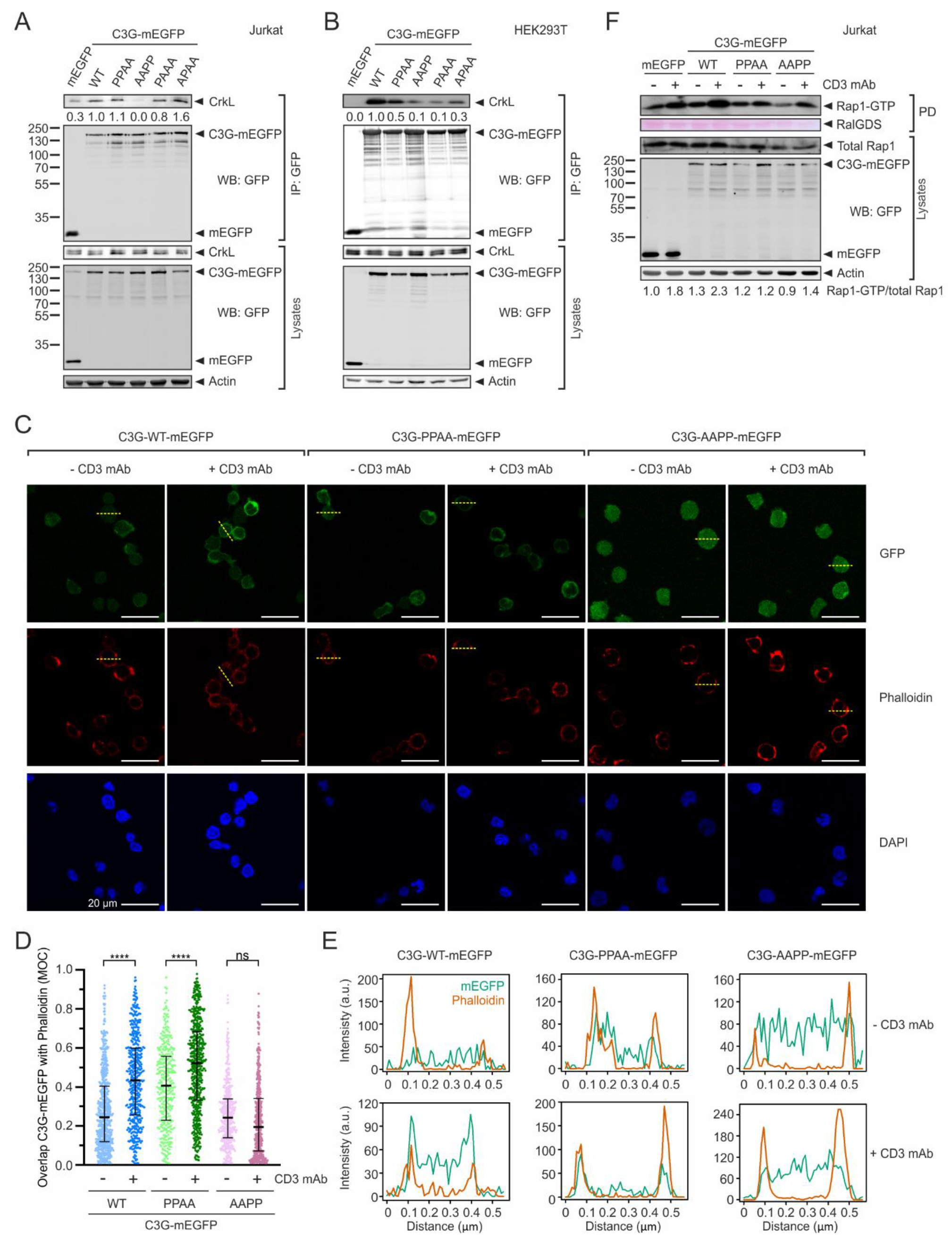
Role of the PRMs in the recruitment of C3G to the plasma membrane and the activation of Rap1 in Jurkat cells. (**A**) Analysis by co-immunoprecipitation (coIP) in Jurkat cells of the interaction between stably expressed exogenous C3G-mEGFP variants and endogenous CrkL. Proteins were immunoprecipitated with affinity resin against GFP. CrkL and mEGFP-tagged proteins were detected in cell lysates and in the IP by western blot (WB). (**B**) Analysis of the interaction between C3G-mEGFP and CrkL in HEK293T cells. mEGFP-tagged C3G WT and mutants were transiently expressed and the interaction with endogenous CrkL was analyzed by coIP as in A. (**C**) Imaging of C3G-mEGFP (upper panels), and cortical actin (stained with phalloidin-iFluor 647, middle panels) in Jurkat cells expressing C3G wild type or the indicated mutants. Nuclei were stained with DAPI. Cells were plated on coverslips treated with poly-L-lysine and were unstimulated (-anti-CD3) or stimulated with OKT-3 antibody against CD3. Representative fields are shown, in which the median of the Manders’ overlap coefficient (MOC) for the cells in each field is similar to the value observed for the total of cells analyzed. Scale bars, 20 μm. (**D**) Quantification of the co-localization of C3G-mEGFP with phalloidin-stained actin in confocal microscopy images of Jurkat cells as shown in C. MOC values are shown as scatter plots (from left to right, n = 637, 466, 346, 565, 423, and 459 cells). Middle bars mark the median and whiskers are the 25th and 75th percentiles. Statistical analysis were done with non-parametric Kruskal-Wallis and Dunn’s multiple comparisons tests (**** *P* < 0.0001, n.s. *P* > 0.05). (**E**) Fluorescence intensity of C3G-mEGFP and phalloidin staining along the yellow dashed lines in panel C, which run across representative cells. (**F**) Analysis of Rap1 activation in Jurkat cells stably expressing C3G-mEGFP WT and mutants, or isolated mEGFP as control, before and after stimulation with antibody against CD3.

We also analyzed the interaction between C3G-mEGFP and CrkL in HEK293T cells (Fig. 5B). HEK293T cells were transiently transfected with constructs coding for wild type and PRM-mutants of C3G-mEGFP, or mEGFP alone as a control, and the interaction was assayed by coIP as above. As observed in Jurkat cells, CrkL interacts strongly with wild type C3G-mEGFP and the PPAA mutant. CrkL coIP less with C3G-AAPP and the single-site mutants C3G-PAAA and C3G-APAA.

We were also interested in understanding the contribution of the interaction of Crk proteins with the two distinct pairs of PRMs of C3G during the activation of Rap1 in cells. C3G is required for the activation of Rap1 in Jurkat cells upon ligation of the T cell receptor (TCR), in a process that involves the recruitment of CrkL/C3G and CrkII/C3G complexes to the plasma membrane [22, 29]. We used Jurkat cells expressing wild type C3G-mEGFP or the mutants containing only wild type P1-P2 (C3G-PPAA) or P3-P4 (C3G-AAPP), to analyze by confocal microscopy the potential translocation of C3G-mEGFP to the cell periphery upon ligation of TCR/CD3 complexes using an antibody against CD3 (Fig. 5C-E). We used phalloidin staining of cortical actin as a marker for the plasma membrane at the resolution of conventional confocal microscopy [30]. Co-localization of wild type C3G-mEGFP and C3G-PPAA-mEGFP with cortical actin increased significantly upon CD3 ligation, suggesting that they were recruited to the plasma membrane. In contrast, there were no differences in the co-localization of C3G-AAPP-mEGFP with cortical actin before and after CD3 ligation.

Next, we analyzed the activation of Rap1 upon ligation of TCR in the Jurkat cells expressing C3G-mEGFP, wild type and PRM mutants (Fig. 5F). These cells express endogenous C3G, which resulted in the activation of Rap1 after CD3 ligation in the presence of mEGFP alone. Nonetheless, cells expressing wild type C3G-mEGFP showed a stronger activation of Rap1, revealing a contribution of the exogenous C3G. Rap1 was not activated upon CD3 ligation in cells expressing C3G-PPAA-mEGFP. This can be interpreted as a dominant negative effect of C3G-PPAA-mEGFP that is not activated by CrkL, but would bind to CrkL reducing its access to the endogenous C3G, which in turn cannot be activated. Finally, in Jurkat cells expressing C3G-AAPP-mEGFP the levels of active Rap1 upon stimulation were similar to those observed in the cells expressing mEGFP, in agreement with the inability of this mutant to be recruited to signaling sites. In summary, C3G-mediated activation of Rap1 in Jurkat cells requires the PRM sites necessary for recruitment of C3G to the plasma membrane (P1 and P2) and those necessary for the direct stimulation of the GEF activity (P3 and P4), likely through the interaction with Crk adaptor proteins.

## Discussion

Understanding the specific mechanisms of autoregulation and activation of different GEFs is essential to comprehending the spatiotemporal regulation of small GTPases. This is the case of Rap1 proteins, which are activated in the same cells by multiple stimuli that engage different GEFs. For example, in platelets Rap1b responds to low thrombin levels through CalDAG-GEF I (RasGRP2), while C3G is the main Rap1GEF at high thrombin stimulation [31-34]. We have elucidated the activation mechanisms of C3G, which improves the understanding of Rap1 signaling.

### Crk proteins stabilize an active state of C3G

We propose that C3G exists in an equilibrium between an inactive or closed state, where the Cdc25HD is blocked, and an active or open conformation where the AIR is dissociated from the Cdc25HD. In resting C3G, this equilibrium would be mostly shifted towards the inactive conformation. The presence of a small fraction of active C3G prior to stimulation is supported by the residual GEF activity of unphosphorylated wild type C3G. A similar equilibrium between inactive and active states has been proposed for other Cdc25H GEFs such as Epac proteins [35].

Binding of Crk proteins to the P3 is the driving event for C3G activation. Access to the P3 is linked to the activation state of C3G. The P3 is adjacent to the AIR-CBR and it is occluded in the inactive state by the AIR-CBR/Cdc25HD interaction. The low stoichiometry of the CrkL binding to P3 in resting C3G could represent the small fraction of active C3G in the conformational equilibrium. The P3 is exposed by the activating mutation Y554H that disrupts binding of the AIR-CBR to the Cdc25HD. Binding of CrkL to the P3 displaces the interaction between the AIR and Cdc25HD when produced as independent proteins [8]. In summary, we propose that binding of Crk proteins to the P3 stabilizes an open conformation and shifts the conformational equilibrium of C3G. Binding of CrkL to the P4 contributes to the optimal activation likely by further stabilizing the active state, but only after CrkL is bound to the P3. In the case of unphosphorylated C3G, stabilization of an active state requires high saturation levels of binding of Crk proteins.

### Role of Src-mediated phosphorylation of C3G during activation

Tyrosine phosphorylation of C3G by Src causes multiple effects that together favor the activation of C3G by Crk proteins. Phosphorylation of C3G does not disrupt the interaction of the AIR-CBR with the Cdc25HD [8] and does not expose the P3 for binding of CrkL. Yet, pC3G has slightly but significantly higher GEF activity compared to unphosphorylated C3G. This may be explained if phosphorylation disturbs the interaction between the AIR-inhibitory tail (AIR-IT) and the Cdc25HD, favoring a partially active state, without affecting the binding of the AIR-CBR to the Cdc25HD.

Based on the comparative activation of C3G and pC3G by CrkL and its fragments, phosphorylation of C3G favors activation by Crk proteins in two ways. First, pC3G is more susceptible to activation by the CrkL-SH3N domain than unphosphorylated C3G, and this effect is independent of the contributions of the SH2 and SH3C domains. Second, phosphorylation creates one or more binding sites for the SH2 domain of CrkL, and this interaction is essential for the efficient activation of pC3G. Full-length CrkL and its isolated SH2 domain bind to a consensus phospho-tyrosine peptide with *k*_d_ values of 23 μM and 7 μM, respectively [36]. Our data indicates that the CrkL-SH2 domain binds to pC3G with lower affinity, suggesting that the SH2 interacts with phospho-tyrosine sites that are not part of a canonical pY-x-x-P motif. Future work would be required to map the phosphorylation site or sites in C3G recognized by the SH2 domain of Crk proteins. The weak interaction between CrkL-SH2 and pC3G also suggests that the interacting SH2 domain belongs to a CrkL molecule that also binds to C3G through the SH3N domain. Initial binding of the SH3N to a PRM of C3G would facilitate the interaction of the companion SH2 domain. The Crk molecule(s) participating in the dual interaction through the SH3N and SH2 domains would not link C3G to other tyrosine phosphorylated proteins. Therefore, the direct activation of pC3G by CrkL and CrkII is, to the best of our knowledge, the first example of Crk proteins signaling through an adaptor-independent mechanism.

CrkL stimulates the GEF activity of pC3G to similar levels as those caused by point mutations that disrupt the AIR/Cdc25HD autoinhibitory interaction, such as Y554R and Y554H. This suggests that CrkL binding to pC3G displaces the AIR/Cdc25HD autoinhibitory interaction and efficiently shifts the conformation equilibrium of C3G, stabilizing an open and active state.

### Model for C3G activation

We propose the following multi-step mechanism for the regulation and physiological activation of C3G (Fig. 6). Prior to stimulation, C3G is diffusely located in the cytoplasm where it interacts with Crk proteins through the P1 and P2 sites. We have shown that activation of unphosphorylated C3G occurs at concentrations of Crk proteins that are more than 10-times higher than those required for binding (AC_50_ ∼10 x *k*_d_). This uncoupling between Crk binding and Crk-mediated activation of C3G, together with the likely relatively low concentration of Crk proteins in the cytoplasm, provides a mechanism for the existence of inactive Crk/C3G complexes. The preformed inactive Crk/C3G complexes probably contribute to a fast response during activation.

**Fig. 6.**
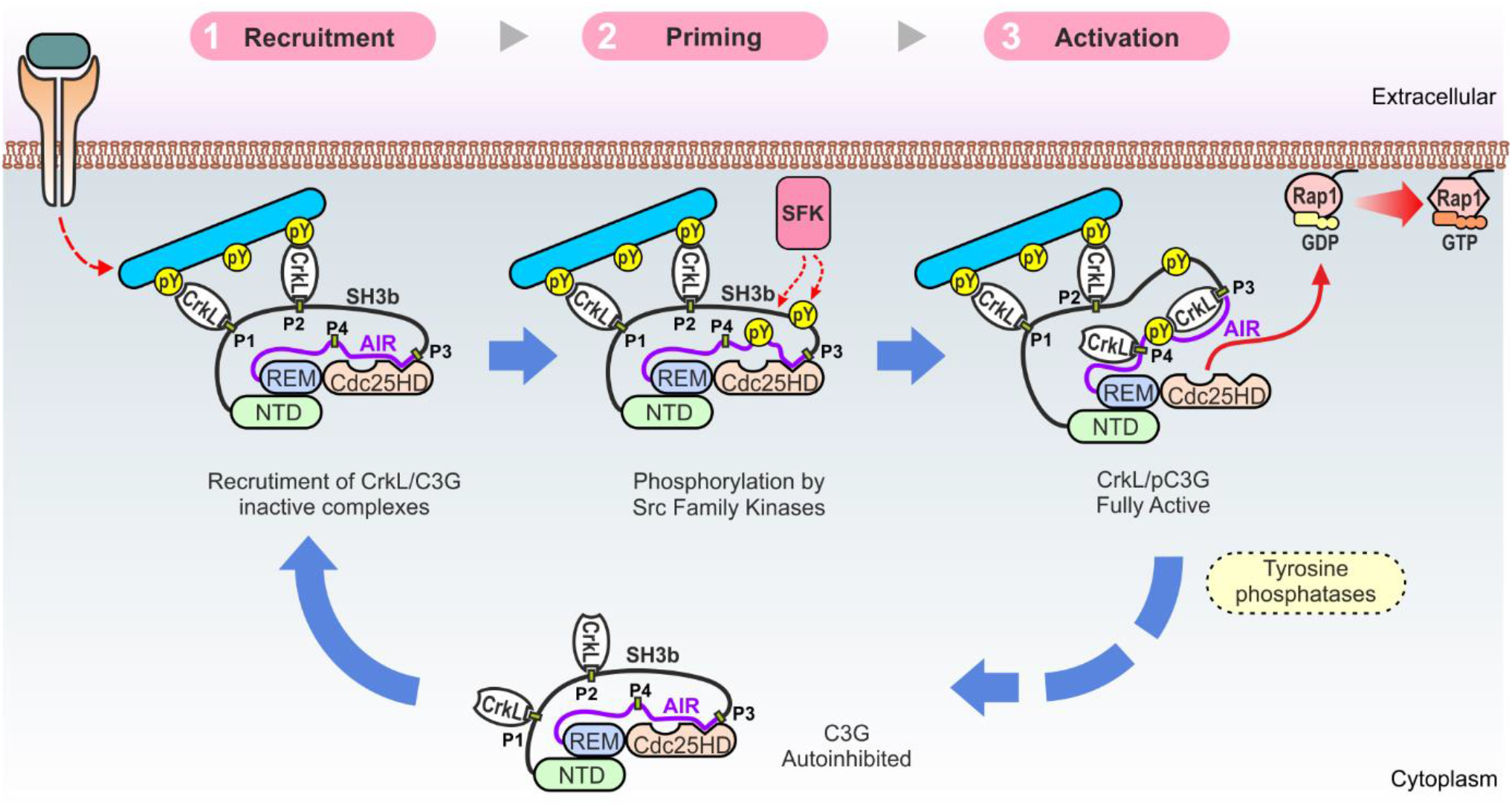
Model for the activation of C3G. Activation of C3G is a three-step process. Initial recruitment relies on the interaction of CrkL with the P1 and P2 sites. Translocation to the plasma membrane is independent of the activation of C3G. The second step is the priming of C3G by phosphorylation in tyrosine residues by Src-family kinases (SFKs) or other kinases such as Abl. Phosphorylation alone is insufficient for activation. The third step is the actual activation, which occurs when CrkL interacts with the P3 and P4 sites in phosphorylated C3G. CrkL stabilizes an open state of pC3G in which the autoinhibitory interaction between the AIR and Cdc25HD is prevented, unleashing the GEF activity of the Cdc25HD. Activation is likely to be reverted, among other mechanisms, through the action of tyrosine phosphatases, which would result in the dissociation of CrkL from the P3 activation site and the diffusion of inactive CrkL/C3G complexes to the cytoplasm.

C3G activation is triggered by signals that stimulate tyrosine kinases at the plasma membrane. In a first step, Crk molecules, bound through the SH3N to mainly the P1 and P2 sites of C3G, bind with the SH2 domain to tyrosine phosphorylated proteins such as p130Cas or paxillin [37, 38]. This results in the translocation of the Crk/C3G complexes to the signaling sites. In this stage, Crk proteins act as canonical adaptors linking C3G to activated phospho-proteins. In a second step, C3G is phosphorylated at tyrosine residues at the plasma membrane, which alone is insufficient for efficient activation. The main role of phosphorylation is the priming of C3G to facilitate its activation by Crk proteins in the final third step. On the one hand, phosphorylation sensitizes C3G reducing the concentrations of Crk proteins required for activation, as reflected by a 10-fold reduction of the CrkL AC_50_ between unphosphorylated and Src-phosphorylated C3G. Phosphorylation of C3G couples binding of CrkL to the activation; that is, the AC_50_ now matches the *k*_d_. On the other hand, C3G phosphorylation increases 3-times the maximal activity induced by CrkL and CrkII with respect to unphosphorylated C3G. Binding of Crk proteins to C3G at signaling sites is likely to be favored by the recruitment of additional molecules of CrkL or CrkII to the phospho-tyrosine sites. Finally, the GEF activity of C3G would be later turned off by the action of tyrosine phosphatases. For example, C3G is dephosphorylated by the T cell protein tyrosine phosphatase TC-PTP (PTPN2) [39], and by the tyrosine phosphatase Shp2 (PTPN11) that inhibits the GEF activity of C3G in platelets [33]. C3G dephosphorylation would result in the rapid dissociation of CrkL, as suggested by the highly dynamic character of the CrkL/C3G interaction, and would lead to the diffusion of inactive C3G in the cytoplasm.

Efficient activation of C3G requires the convergence of tyrosine phosphorylation of C3G and binding of Crk proteins. This two-factor mechanism allows for a tight spatiotemporal regulation of C3G signaling, which would only be activated when and where both events concur. Notably, two key steps in C3G activation, recruitment to signaling sites and unlocking of the GEF activity, are mediated by the same Crk proteins. We have shown that these two events are segregated along the sequence of C3G and involve binding of Crk to different sites. The four Crk-binding sites of C3G are chemically equivalent; that is, CrkL binds to the four PRMs with similar affinity. Yet, these sites are functionally non-equivalent and are not equally accessible in full-length C3G. Binding of Crk proteins to the P1 and P2 is linked to the translocation of C3G, while binding to the second pair of PRMs is necessary for the stimulation of the GEF activity. Therefore, we refer to the P1 and P2 as recruitment-sites and to the P3 and P4 as activation-sites. Our findings highlight the importance of analyzing the interactions of PRMs and other short linear motifs in the context of full-length proteins.

### Implications of C3G activation mechanism in physiological signaling and diseases

The more efficient activation of pC3G by CrkL than by CrkII might contribute to differences in Rap1 signaling by the two Crk proteins in vivo. For example, Rap1 activation induced by chemokines in T cells plays a key role during trans-endothelial migration or diapedesis [40]. Knockout of CrkL causes stronger defects in diapedesis of T cells than depletion of CrkII [41]. This suggests that Rap1 activation in response to chemokines is more dependent on CrkL than on CrkII, in agreement with our findings in vitro. Yet, other factors may also contribute to the differential signaling by CrkII and CrkL including expression levels and regulatory mechanisms.

The levels of Crk proteins are increased in a variety of human cancers and is linked to poor prognosis [42]. Ectopic overexpression of CrkL or CrkII stimulates the C3G-dependent activation of Rap1 [8, 43], suggesting that increased abundance of Crk proteins in malignant cells may cause abnormal activation of C3G-Rap1. The proposed model of C3G-Rap1 activation implies that, in scenarios of elevated concentrations of Crk proteins, the activation of C3G would not require, or would depend less on, tyrosine phosphorylation of C3G. In addition, high levels of Crk proteins may induce the abnormal targeting of C3G to sites with low levels of tyrosine phosphorylation that would be insufficient to recruit C3G under physiological conditions. This is supported by the increase in phosphorylation of C3G induced by overexpression of Crk [18].

In conclusion, the detailed mechanistic description of the activation of C3G presented here helps to comprehend physiological CrkL-C3G-Rap1 signaling and paves the way to analyze alterations in this pathway linked to diseases.

## Materials and methods

### DNA constructs and site directed mutagenesis

Constructs for the expression of C3G in bacteria are listed in Table S6. The cDNA of human C3G (isoform a, residues 4-1077, UniProt Q13905-1) was cloned into a modified vector pETEV15b [44] that codes for an N-terminal poly-His tag, the Halo protein, and a site recognized by the tobacco etch virus (TEV) protease (pETEV15b-His-Halo-TEV). C3G cDNA was amplified by the polymerase chain reaction (PCR) using as template a C3G construct in a derivative of the vector pGEX-4T3 [8], in which the internal NdeI was destroyed by a silent mutation introduced by PCR-based site directed mutagenesis. The primers added NdeI and BamHI sites at the 5’ and 3’ ends, respectively, and the amplified cDNA was cloned into those sites in the pETEV15b-His-Halo-TEV vector. The Y554H point mutation and the Pro-to-Ala mutations in the PRMs were introduced by site directed mutagenesis. Primers used to create C3G constructs and to introduce mutations are shown in Table S7. Correctness of these and all other constructs was confirmed by DNA sequencing.

Constructs of C3G for expression in mammalian cells are listed in Table S8. C3G in the lentiviral vector pLenti-C-mEGFP-IRES-BSD has been previously described [8]. C3G mutants C3G-PPAA, C3G-AAPP, C3G-PAAA, and C3G-APAA in pLenti-C-mEGFP-IRES-BSD were created by PCR-amplification of the cDNA from the corresponding bacterial expression constructs. The primers added BamHI and NotI sites at the 5’ and 3’ ends and the reverse primer did not include a stop codon. The amplified cDNAs were ligated into pLenti-C-mEGFP-IRES-BSD using BamHI and NotI sites. C3G in the vector pEF1-mEGFP has been described [8], the aforementioned PRM mutants of C3G in the lentiviral expression vector were also cloned in pEF1-mEGFP using the same BamHI and NotI restriction sites as above.

Constructs of human CrkL (UniProt P46109) and CrkII (UniProt P46108-1) for bacterial expression are shown in Table S9. A construct of human CrkL in the vector pETEV15b has already been described [8]. The cDNAs coding for the CrkL fragments SH3N (residues 125-182) and SH2-SH3N (1-182) were amplified by PCR adding NdeI and BamHI sites at the 5’ and 3’ ends. The cDNA of CrkL SH3N-SH3C (125-303) was PCR-amplified similarly, but adding a BglII site at the 3’ end. The fragments were cloned into the NdeI and BamHI sites of pETEV15b. The cDNA of full-length CrkL and the SH3N domain (125-182) were similarly cloned in a variant of pGEX-4T3-TEV that has NdeI and BamHI sites for cloning [8]. The cDNA of CrkL SH3N (111-204) was PCR-amplified and cloned in pGEX-4T3 using SalI and NotI sites. Point mutations R39K and W160S were introduced in CrkL constructs by site directed mutagenesis. The cDNA of human CrkII was amplified by PCR adding NdeI and BamHI sites and was cloned into pETEV15b using these sites. The chimeric constructs CrkL-II-L (residues CrkL 1-124, CrkII 134-191, and CrkL 183-303) and CrkII-L-II (residues CrkII 1-133, CrkL 125-182, and CrkII 191-304) were created by overlap extension PCR and were cloned in pETEV15b using NdeI and BamHI in the PCR products and the vector. Prior to amplifying the cDNA of CrkL 183-303 the internal BamHI site was destroyed introducing a silent mutation by site directed mutagenesis. Oligonucleotides used for creating the CrkL and CrkII constructs are shown in Table S10.

The cDNA of the nanobody abGFP4 recognizing GFP (a gift from Brett Collins; Addgene plasmid 49172) [45] was subcloned into a derivative of pET22b vector that lacks the pelB sequence and codes for a C-terminal poly-His tag followed by a cysteine. The nanobody cDNA was PCR-amplified with primers that added NcoI and XhoI sites at the 5’ and 3’ ends (Table S11) and was cloned into those sites of the vector.

### Expression and purification of recombinant proteins

Proteins were produced in *E. coli* strain BL21(DE3) grown in Terrific Broth medium with 100 µg/ml ampicillin. Expression was induced at 15°C overnight with 0.2 mM isopropyl-β-D-thiogalactopyranoside. Full-length C3G was produced as a fusion protein with N-terminal poly-His and Halo tag, the latter increased the expression levels. C3G was first purified by immobilized metal affinity chromatography (IMAC) using Ni^2+^-chelating sepharose (Cytiva) or agarose (Agarose Bead Technologies, ABT) columns as described [46]. Next, the poly-His and Halo tags were cleaved by digestion with TEV protease and were removed in a second reversed IMAC. The fractions containing C3G were concentrated using Amicon ultrafiltration cells and centrifugal filters (Millipore) and were further purified by size exclusion chromatography (SEC) in a Superdex 200 (1 × 30 cm) column equilibrated in 20 mM Tris-HCl, 300 mM NaCl, pH 7.5. Finally, the samples were concentrated by ultrafiltration, flashed freeze in liquid nitrogen, and stored at -80°C until used. C3G proteins were phosphorylated in vitro with the recombinant kinase domain of Src as described [8]. After the reaction, phosphorylated C3G was purified by SEC using a Superdex 200 (1 × 30 cm) column equilibrated in 20 mM Tris-HCl, 300 mM NaCl, pH 7.5.

His-tagged CrkL proteins were purified in a similar manner. GST-CrkL-SH3N was purified by affinity chromatography using glutathione agarose columns (ABT) as described [8].

The abGFP4 nanobody against GFP was first purified by IMAC as for the C3G proteins. Next, it was concentrated by ultrafiltration and was subjected to SEC using a HiPrep Sephacryl S-200 HR (2.6 × 60 cm) column (Cytiva) equilibrated in 20 mM Tris-HCl, 150 mM NaCl, pH 7.5. Finally, the nanobody was concentrated by ultrafiltration and stored at -80°C.

### Fluorescence anisotropy-based binding assay

Peptides corresponding to the P1 (residues 280-293), P2 (450-463), P3 (537-550), and P4 (605-618) carrying an

N-terminal fluorescein probe, were custom synthesized (Thermo Scientific). Binding of CrkL to these peptides was analyzed by fluorescence anisotropy as described [47]. Briefly, peptides at 0.2 µM in 20 mM Tris-HCl, 150 mM NaCl, pH 7.5, 0.1 mg/ml bovine serum albumin (BSA), were titrated with CrkL. The fluorescence anisotropy was measured in an Ultra Evolution plate reader (Tecan) using 485 nm and 535 nm excitation and emission filters. The apparent dissociation constant (*k*_d_) was determined by nonlinear square fitting of a 1:1 binding model using SigmaPlot (Systat Software Inc.).

### Isothermal titration calorimetry (ITC)

ITC experiments were performed using a MicroCal VP-ITC calorimeter (Malvern Pananalytical) at 25°C. The working buffer was either 20 mM Tris-HCl, 300 mM NaCl, pH 7.5 or 20 mM Na-phosphate, 300 mM NaCl, pH 7.5. Solutions of C3G proteins between 5 and 20 µM in the cell were titrated with CrkL or CrkII proteins between 100 and 200 µM. Heat signals were integrated with the program NITPIC [48]. Data of single experiments were initially analyzed with the program MicroCal Origin ITC module (Malvern, version 7.0) using an independent-site association model, which fits the association constant (*k*_a_ = 1/*k*_d_) and the relative stoichiometry (N). Alternatively, ITC experiments were analyzed individually or globally with the software SEDPHAT [49] using two hetero-association models. Interactions of CrkL with wild type C3G were analyzed using a model with three symmetric binding sites; in these cases, microscopic *k*_d_ values were reported. The binding of CrkL to C3G mutants displaying a single PRM was analyzed using a 1:1 binding model. SEDPHAT does not fit non-integral binding sites; instead, the incompetent fraction of C3G was fitted and the fraction of competent or active binding sites was estimated as the complementary value.

### Sedimentation velocity (SV)

SV experiments were performed in an Optima XL-I analytical ultracentrifuge (Beckman-Coulter Inc.) using an An-50Ti rotor and 12 mm optical pass double sector cells. Samples of C3G and CrkL alone, or C3G titrated with several concentrations of CrkL in 20 mM Tris-HCl, 300 mM NaCl, pH 7.5 were subjected to centrifugation at 48000 rpm (167700 *g*) at 20°C and sedimentation was followed by interference and absorbance at 280 nm. Differential sedimentation coefficient distributions were calculated by least-squares boundary modeling of sedimentation velocity data using the continuous distribution c(*s*) Lamm equation model as implemented by SEDFIT [50]. The experimental s values were corrected to standard conditions of water at 20ºC with SEDNTERP [51] to obtain the corresponding standard s values (*s*_20,w_) using the partial specific volumes (v_bar_) of C3G (0.733 cm^3^/g) and CrkL (0.728 cm^3^/g) calculated from their sequences or, in the case of C3G-CrkL mixtures, calculated as an average of the values of the individual species (0.730 cm^3^/g). The concentration dependent changes in the shapes of the reaction boundaries were modeled by isotherms based on the Gilbert-Jenkins theory that provides a robust approach to exploit the bimodal structure of the reaction boundary [52]. Integrated weight-average sedimentation coefficients, s-values of the observed fastest boundaries and signal amplitudes for the undisturbed and reaction boundaries, were assembled into isotherms and globally analyzed through a binding model with three symmetric binding sites as implemented in SEDPHAT [53]. In a step beyond, these SV isotherms together with those obtained from ITC experiments were globally analyzed through a global multi-method analysis using the same three symmetric binding sites model in SEDPHAT.

### Dynamic light scattering (DLS)

Experimental diffusion coefficients (D) were obtained by DLS using a DynaPro MS/X instrument (Protein Solutions) and a 90° light scattering cuvette. Samples were analyzed by DLS in the same experimental conditions used in the SV experiments. The apparent molar masses (M) of C3G and CrkL were calculated using measured values of the sedimentation coefficient *s* and the diffusion coefficient D according to the Svedberg equation [54]:

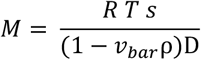

Where T, R, ν_bar_ and ρ are the absolute temperature, the universal gas constant, the partial specific volume of the protein and the density of the solution, respectively.

### Nucleotide exchange activity assays

Nucleotide exchange activity of C3G toward Rap1b was measured in vitro using a Rap1b bound to the GDP fluorescent analogue 2’-deoxy-3’-O-(N’-methylanthraniloyl)guanosine-5’-O-diphosphate (mant-dGDP) as described [8]. Briefly, Rap1:mant-dGDP (200 nM) in 50 mM Tris-HCl, 150 mM NaCl, 10 mM MgCl_2_, pH 7.5 was incubated with C3G proteins in the absence or presence of CrkL or CrkII proteins. The exchange reactions were done at 25ºC and were initiated by adding 40 µM GDP. The fluorescence of mant-dGDP was measured in a FluoroMax-3 fluorometer using excitation light of 370 nm (1 nm bandwidth) and the emission at 430 nm (10 nm bandwidth). The time-dependent decay of the fluorescence intensity was analyzed by fitting a single exponential decay model. The fitting yielded the apparent dissociation rate constant (*k*_obs_), the initial intensity before starting the reaction and the intensity of the fully dissociated mant-dGDP. For representation, data were normalized between 1 (initial intensity) and 0 (final intensity after complete dissociation). The exchange rate constants are reported as specific activity values normalized by the concentration of C3G.

When the exchange rate constants were determined for a range of concentrations of C3G, the dependence of *k*_obs_ with the concentration of C3G was analyzed fitting the following linear equation:

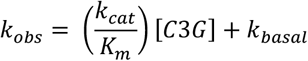

Where, *k*_basal_ is the rate constant of the spontaneous dissociation in the absence of C3G, and *k*_cat_/*K*_m_ is the apparent second order rate constant of a Michaelis–Menten mechanism [55], which represents the nucleotide exchange efficiency of C3G.

### Analysis of dose-response activation of C3G

The dose-dependent activation of C3G by CrkL or CrkII proteins was analyzed by measuring nucleotide dissociation experiments at a fixed concentration of C3G and multiple concentrations of Crk proteins. The variations of the nucleotide exchange rate constants of C3G were analyzed by fitting the following sigmoidal equation:

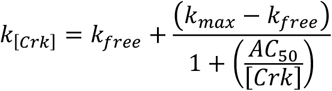

Where, *k*_[Crk]_ is the *k*_obs_ determined at a given concentration of the CrkL or CrkII proteins; *k*_max_ is the maximum value of the rate constant that corresponds to saturated conditions, *k*_free_ is the minimum value of *k*_obs_ that is the activity in the absence of Crk proteins, and AC_50_ is the concentration of Crk proteins that produces the half maximal activation. Analysis was done with SigmaPlot (Systat Software Inc.).

### Antibodies and western blotting

The following primary antibodies were used for western blot, at 1/1000 dilutions unless otherwise specified: mouse monoclonal antibody (mAb) G9 against C3G (sc-393836), rabbit polyclonal antibody (pAb) C-20 against CrkL (sc-319), mouse mAb B-2 against GFP (sc-9996), rabbit pAb 121 against Rap1 (sc-65) were from Santa Cruz Biotechnology, mouse mAb 4G10 against phospho-Tyr (05-321, Millipore), and mouse mAb against β-actin (AC-15, Sigma-Aldrich, used at 1/2000).

The following secondary antibodies were used (1/5000) for western blot: horseradish peroxidase (HRP)-conjugated goat antibody against rabbit IgG (sc-2004, Santa Cruz Biotechnology), HRP-conjugated sheep antibody against mouse IgG (NXA931, Cytiva), goat pAb DyLight 680 against mouse IgG, goat pAb DyLight 680 against rabbit IgG, goat pAb DyLight 800 against mouse IgG, and goat pAb DyLight 800 against rabbit IgG (Thermo Fisher Scientific). Signals from secondary antibodies were recorded and quantified as described [8].

### Affinity pull-down assays

Pull-down (PD) assays were done using purified GST-fusion CrkL proteins and C3G proteins expressed in *E. coli*. Typically, 30 µg of GST-CrkL (full-length or SH3N), or GST alone as a control, were mixed with 30 µg of C3G in 20 mM Tris-HCl, 150 mM NaCl, pH 7.5, 0.01% BSA, and 20 µl of glutathione-agarose resin (ABT). The samples were incubated for 30 min at 4°C. The resin was washed four times with the previous buffer supplemented with 0.1% Triton X-100, and the proteins were extracted by adding Laemmli sample buffer. Proteins in the input and PD samples were detected either by coomassie staining or by western blot with antibodies that recognize C3G or phospho-Tyr.

### Cell lines

HEK293T cells were grown in Dulbecco’s modified Eagle’s medium (DMEM; Sigma-Aldrich) with 10% fetal bovine serum (FBS; Life Technologies), 100 U/ml penicillin, and 100 µg/ml streptomycin (Life Technologies). HEK293T cells were transfected with plasmids using polyethylenimine (PEI; Polysciences Inc.) at PEI:DNA ratio 2.5:1. Jurkat cells were grown in RPMI 1640 medium (Sigma-Aldrich) with 10% FBS, 100 U/ml penicillin, and 100 µg/ml streptomycin. All cells were grown at 37°C and 5% CO_2_.

### Lentiviral production and cell transduction

Lentiviral particles were produced in HEK293T cells. Constructs of C3G in pLenti-C-mEGFP-IRES-BSD lentiviral transfer vector [8], or the empty vector expressing mEGFP, were co-transfected with plasmids pMD2.G (provided by D. Trono; Addgene no. 12259) and pCMV-deltaR8.91 (Lifescience market). Jurkat cells were infected with the recombinant viruses in the presence of 8 µg/ml polybrene (hexadimethrine bromide, Sigma-Aldrich) for 15 hours at 37°C. Cells expressing C3G-mEGFP or mEGFP were selected with 10 μg/ml blasticidin (InvivoGen) for 3 weeks. Afterwards, cells expressing mEGFP were separated by fluorescence-activated cell sorting.

### Co-immunoprecipitation (coIP) assays

Jurkat cells expressing C3G-EGFP, wild type or mutants forms, were used for coIP assays. Typically, ∼30 × 10^6^ cells were lysed in 300 µl of lysis buffer consisting of 20 mM Tris-HCl (pH 7.5), 150 mM NaCl, 1 mM EDTA, 0.5% Triton X-100, 1 mM sodium orthovanadate, 25 mM NaF, 1 mM phenylmethylsulfonyl fluoride (PMSF), and 1x protease inhibitor cocktail (cOmplete, Roche). Cell debris were removed by centrifugation (16000 *g*, 20 min, 4°C) and the protein concentration in the supernatant was determined by Bradford assay. Clarified lysates containing 1 mg of total protein were incubated with 20 µl of 50% slurry of abGFP4 nanobody resin against GFP (see below) and the volume was completed to 500 µl with lysis buffer. The reaction samples were incubated overnight at 4°C with gentle mixing. The resins were washed three times with lysis buffer. Bound proteins were extracted with Laemmli sample buffer and were boiled. Samples of the cell lysates and the IPs were analyzed by western blot.

HEK293T cells transiently transfected with constructs in the pEF1-mEGFP vector coding for C3G-mEGFP or mEGFP alone were lysed 48 hours after transfection. Cells from a 10 cm diameter plate were lysed in 500 μl of lysis buffer and the coIP was done as for the Jurkat cells (see above).

The recombinant abGFP4 nanobody against GFP with a C-terminal cysteine was covalently coupled to SulfoLink resin (Thermo Scientific) consisting of cross-linked agarose beads with iodoacetyl groups. The nanobody in 20 mM Tris-HCl, 150 mM NaCl, pH 7.5 was reduced with 5 mM tris(2-carboxyethyl)phosphine (TCEP) for 1 hour at room temperature. Next, it was mixed with the resin equilibrated in coupling buffer (50 mM Tris-HCl, 5 mM EDTA, pH 8.5) in a ratio of 4 µg of nanobody per µl of resin, and the reaction was incubated for 1 hour at room temperature. The unbound sample was removed and the resin was washed four times with coupling buffer. Unreacted groups in the resin were blocked by incubation with 50 mM L-cysteine in coupling buffer (1 hour at room temperature). The resin was washed three times with 1 M NaCl, was equilibrated in 20 mM Tris-HCl, 150 mM NaCl, pH 7.5, 50% glycerol, and stored at -20°C until used.

### Confocal fluorescence microscopy

Jurkat cells were adhered to coverslips treated with poly-L-lysine during 30 min at 37°C in RPMI. Cells were stimulated with 5 μg/ml of antibody OKT-3 recognizing CD3 (eBiosciences) for 30 min at 37°C. Cells were fixed in 1% paraformaldehyde in phosphate buffer saline (PBS), permeabilized with 0.2% Triton X-100 in PBS for 10 min and stained with phalloidin-iFluor 647 (ab176759, Abcam) for 20 min. Cells were washed three time in PBS and nuclei were stained with 4’,6-diamidino-2-phenylindole (DAPI, Sigma). Coverslips were mounted onto glass slides in Mowiol-DABCO. Cells were observed in a Leica SP5 confocal microscope.

The degree of co-localization between C3G-mEGFP and phalloidin-stained actin in confocal microscopy images was quantified using the Manders’ overlap coefficient (MOC) calculated with the Coloc2 plugin in the Fiji distribution of ImageJ [56].

### Rap1 activation assays in cells

Active Rap1-GTP was detected in Jurkat cells by PD using the fusion protein GST-RalGDS-RBD as described [8] with minor changes. Briefly, 5 × 10^6^ Jurkat cells in RPMI were stimulated with 5 μg/ml of antibody OKT-3 against CD3 (eBiosciences). 30 min after stimulation cells were harvested by centrifugation and were lysed in 200 μl of magnesium lysis buffer (MLB) (25 mM HEPES pH 7.5, 150 mM NaCl, 1% Igepal CA-630, 10 mM MgCl_2_, 1 mM EDTA, 2% glycerol, 1 mM Na_3_VO_4_, 1 mM PMSF, 1x protease inhibitor cocktail (Roche), and 1 mM DTT) in the presence of 30 μg of purified GST-RalGDS-RBD. Samples were clarified by centrifugation (16000 g, 15 min, 4°C). Afterwards, 20 μl of glutathione-agarose resin was added to each sample and it incubated for 10 min at 4°C. The resin was washed three times with 1 ml of MLB. Bound proteins were extracted with SDS-PAGE sample buffer with 10 mM DTT for 12 hours at room temperature. Proteins were analyzed by SDS-PAGE and western blot with an antibody against Rap1 and HRP-conjugates secondary antibody. Total Rap1 was detected in the cell lysates and Rap1-GTP in the PD. Two independent experiments were performed.

### Quantitation and statistical analysis

The normal distribution of data in the groups to be compared was first analyzed using the Shapiro-Wilk test. When data followed a normal distribution (Fig. 3C and 3E) the comparisons of multiple groups of data were done with one-way analysis of variance (ANOVA) with Tukey’s test because variances where equivalent as analyzed with the Brown-Forsythe homogeneity of variance test. For data not following a normal distribution (Fig. 5D), statistical comparisons were done using unpaired non-parametric Kruskal-Wallis and Dunn’s multiple comparisons tests. Differences between groups were considered significant at *P* < 0.05. Significant levels were marked as * *P* < 0.05, ** *P* < 0.01, *** *P* < 0.001, **** *P* < 0.0001. Statistical analyses were done with the program GraphPad Prism 8.

## Data availability

All data needed to evaluate the conclusions in the paper are present in the paper or the Supplementary Materials.

## Acknowledgments

We thank the Biochemical and Biophysical Technologies scientific platforms of i3S (Porto, Portugal) for support during ITC experiments. We thank Brett Collins (University of Queensland, Australia) and Didier Trono (Ecole Polytechnique Fédérale de Lausanne, Switzerland) for providing plasmids. We thank members of the de Pereda and Guerrero laboratories for critical suggestions to the manuscript. JRLO acknowledges support from the Molecular Interactions Facility at the CIB-Margarita Salas CSIC.

## Author contributions

AC, ARB, and JMdP, conceived the study; ARB, AC, and JAM performed the ITC analysis with guidance of SMR; AC, JRLO, and CA performed the SV and DLS experiments and did the global analysis with PS; SdC participated in biochemical analyses, AMV participated in cellular assays, CG participated in the design and interpretation of analyses in cells; AC, ARB, and JMdP wrote the initial drafts for the manuscript and made the figures. All authors contributed and approved the final manuscript.

## References

1. Gotoh T, Hattori S, Nakamura S, Kitayama H, Noda M, Takai Y, Kaibuchi K, Matsui H, Hatase O, Takahashi H. (1995) Identification of Rap1 as a target for the Crk SH3 domain-binding guanine nucleotide-releasing factor C3G. Mol Cell Biol 15(12):6746–53. https://doi.org/10.1128/mcb.15.12.6746.

2. van den Berghe N, Cool RH, Horn G, Wittinghofer A. (1997) Biochemical characterization of C3G: an exchange factor that discriminates between Rap1 and Rap2 and is not inhibited by Rap1A(S17N). Oncogene 15(7):845–50. https://doi.org/10.1038/sj.onc.1201407.

3. Chiang SH, Baumann CA, Kanzaki M, Thurmond DC, Watson RT, Neudauer CL, Macara IG, Pessin JE, Saltiel AR. (2001) Insulin-stimulated GLUT4 translocation requires the CAP-dependent activation of TC10. Nature 410(6831):944–8. https://doi.org/10.1038/35073608.

4. Arai A, Nosaka Y, Kohsaka H, Miyasaka N, Miura O. (1999) CrkL activates integrin-mediated hematopoietic cell adhesion through the guanine nucleotide exchange factor C3G. Blood 93(11):3713–22. https://doi.org/10.1182/blood.V93.11.3713.

5. Radha V, Mitra A, Dayma K, Sasikumar K. (2011) Signalling to actin: role of C3G, a multitasking guanine-nucleotide-exchange factor. Biosci Rep 31(4):231–44. https://doi.org/10.1042/BSR20100094.

6. Martin-Granado V, Ortiz-Rivero S, Carmona R, Gutiérrez-Herrero S, Barrera M, San-Segundo L, Sequera C, Perdiguero P, Lozano F, Martín-Herrero F, González-Porras JR, Muñoz-Chápuli R, Porras A, Guerrero C. (2017) C3G promotes a selective release of angiogenic factors from activated mouse platelets to regulate angiogenesis and tumor metastasis. Oncotarget 8(67):110994–1011. https://doi.org/10.18632/oncotarget.22339.

7. Hogan C, Serpente N, Cogram P, Hosking CR, Bialucha CU, Feller SM, Braga VM, Birchmeier W, Fujita Y. (2004) Rap1 regulates the formation of E-cadherin-based cell-cell contacts. Mol Cell Biol 24(15):6690–700. https://doi.org/10.1128/MCB.24.15.6690-6700.2004.

8. Carabias A, Gomez-Hernandez M, de Cima S, Rodriguez-Blazquez A, Moran-Vaquero A, Gonzalez-Saenz P, Guerrero C, de Pereda JM. (2020) Mechanisms of autoregulation of C3G, activator of the GTPase Rap1, and its catalytic deregulation in lymphomas. Sci Signal 13(647):eabb7075. https://doi.org/10.1126/scisignal.abb7075.

9. Green MR, Gentles AJ, Nair RV, Irish JM, Kihira S, Liu CL, Kela I, Hopmans ES, Myklebust JH, Ji H, Plevritis SK, Levy R, Alizadeh AA. (2013) Hierarchy in somatic mutations arising during genomic evolution and progression of follicular lymphoma. Blood 121(9):1604–11. https://doi.org/10.1182/blood-2012-09-457283.

10. Morin RD, Mendez-Lago M, Mungall AJ, Goya R, Mungall KL, Corbett RD, Johnson NA, Severson TM, Chiu R, Field M, Jackman S, Krzywinski M, Scott DW, Trinh DL, Tamura-Wells J, Li S, Firme MR, Rogic S, Griffith M, Chan S, Yakovenko O, Meyer IM, Zhao EY, Smailus D, Moksa M, Chittaranjan S, Rimsza L, Brooks-Wilson A, Spinelli JJ, Ben-Neriah S, Meissner B, Woolcock B, Boyle M, McDonald H, Tam A, Zhao Y, Delaney A, Zeng T, Tse K, Butterfield Y, Birol I, Holt R, Schein J, Horsman DE, Moore R, Jones SJ, Connors JM, Hirst M, Gascoyne RD, Marra MA. (2011) Frequent mutation of histone-modifying genes in non-Hodgkin lymphoma. Nature 476(7360):298–303. https://doi.org/10.1038/nature10351.

11. Birge RB, Kalodimos C, Inagaki F, Tanaka S. (2009) Crk and CrkL adaptor proteins: networks for physiological and pathological signaling. Cell Commun Signal 7:13. https://doi.org/10.1186/1478-811X-7-13.

12. Birge RB, Fajardo JE, Reichman C, Shoelson SE, Songyang Z, Cantley LC, Hanafusa H. (1993) Identification and characterization of a high-affinity interaction between v-Crk and tyrosine-phosphorylated paxillin in CT10-transformed fibroblasts. Mol Cell Biol 13(8):4648–56. https://doi.org/10.1128/mcb.13.8.4648-4656.1993.

13. Knudsen BS, Feller SM, Hanafusa H. (1994) Four proline-rich sequences of the guanine-nucleotide exchange factor C3G bind with unique specificity to the first Src homology 3 domain of Crk. J Biol Chem 269(52):32781–7. https://doi.org/10.1016/S0021-9258(20)30059-4.

14. Muralidharan V, Dutta K, Cho J, Vila-Perello M, Raleigh DP, Cowburn D, Muir TW. (2006) Solution structure and folding characteristics of the C-terminal SH3 domain of c-Crk-II. Biochemistry 45(29):8874–84. https://doi.org/10.1021/bi060590z.

15. Okada S, Pessin JE. (1997) Insulin and epidermal growth factor stimulate a conformational change in Rap1 and dissociation of the CrkII-C3G complex. J Biol Chem 272(45):28179–82. https://doi.org/10.1074/jbc.272.45.28179.

16. Smit L, van der Horst G, Borst J. (1996) Sos, Vav, and C3G participate in B cell receptor-induced signaling pathways and differentially associate with Shc-Grb2, Crk, and Crk-L adaptors. J Biol Chem 271(15):8564–9. https://doi.org/10.1074/jbc.271.15.8564.

17. Feller SM. (2001) Crk family adaptors-signalling complex formation and biological roles. Oncogene 20(44):6348–71. https://doi.org/10.1038/sj.onc.1204779.

18. Ichiba T, Hashimoto Y, Nakaya M, Kuraishi Y, Tanaka S, Kurata T, Mochizuki N, Matsuda M. (1999) Activation of C3G guanine nucleotide exchange factor for Rap1 by phosphorylation of tyrosine 504. J Biol Chem 274(20):14376–81. https://doi.org/10.1074/jbc.274.20.14376.

19. Mitra A, Radha V. (2010) F-actin-binding domain of c-Abl regulates localized phosphorylation of C3G: role of C3G in c-Abl-mediated cell death. Oncogene 29(32):4528–42. https://doi.org/10.1038/onc.2010.113.

20. Radha V, Rajanna A, Swarup G. (2004) Phosphorylated guanine nucleotide exchange factor C3G, induced by pervanadate and Src family kinases localizes to the Golgi and subcortical actin cytoskeleton. BMC Cell Biol 5:31. https://doi.org/10.1186/1471-2121-5-31.

21. Shivakrupa R, Radha V, Sudhakar C, Swarup G. (2003) Physical and functional interaction between Hck tyrosine kinase and guanine nucleotide exchange factor C3G results in apoptosis, which is independent of C3G catalytic domain. J Biol Chem 278(52):52188–94. https://doi.org/10.1074/jbc.M310656200.

22. Nolz JC, Nacusi LP, Segovis CM, Medeiros RB, Mitchell JS, Shimizu Y, Billadeau DD. (2008) The WAVE2 complex regulates T cell receptor signaling to integrins via Abl- and CrkL-C3G-mediated activation of Rap1. J Cell Biol 182(6):1231–44. https://doi.org/10.1083/jcb.200801121.

23. Gutierrez-Berzal J, Castellano E, Martin-Encabo S, Gutierrez-Cianca N, Hernandez JM, Santos E, Guerrero C. (2006) Characterization of p87C3G, a novel, truncated C3G isoform that is overexpressed in chronic myeloid leukemia and interacts with Bcr-Abl. Exp Cell Res 312(6):938–48. https://doi.org/10.1016/j.yexcr.2005.12.007.

24. Popovic M. (2013) Regulation and Selectivity of Exchange Factors for G-proteins of the Ras-family. Dissertation, Utrecht University.

25. Knudsen BS, Zheng J, Feller SM, Mayer JP, Burrell SK, Cowburn D, Hanafusa H. (1995) Affinity and specificity requirements for the first Src homology 3 domain of the Crk proteins. EMBO J 14(10):2191–8. https://doi.org/10.1002/j.1460-2075.1995.tb07213.x.

26. Schuck P. (2010) Diffusion of the reaction boundary of rapidly interacting macromolecules in sedimentation velocity. Biophys J 98(11):2741–51. https://doi.org/10.1016/j.bpj.2010.03.004.

27. Wu X, Knudsen B, Feller SM, Zheng J, Sali A, Cowburn D, Hanafusa H, Kuriyan J. (1995) Structural basis for the specific interaction of lysine-containing proline-rich peptides with the N-terminal SH3 domain of c-Crk. Structure 3(2):215–26. https://doi.org/10.1016/s0969-2126(01)00151-4

28. Reedquist KA, Fukazawa T, Panchamoorthy G, Langdon WY, Shoelson SE, Druker BJ, Band H. (1996) Stimulation through the T cell receptor induces Cbl association with Crk proteins and the guanine nucleotide exchange protein C3G. J Biol Chem 271(14):8435–42. https://doi.org/10.1074/jbc.271.14.8435.

29. Azoulay-Alfaguter I, Strazza M, Peled M, Novak HK, Muller J, Dustin ML, Mor A. (2017) The tyrosine phosphatase SHP-1 promotes T cell adhesion by activating the adaptor protein CrkII in the immunological synapse. Sci Signal 10(491). https://doi.org/10.1126/scisignal.aal2880.

30. Clausen MP, Colin-York H, Schneider F, Eggeling C, Fritzsche M. (2017) Dissecting the actin cortex density and membrane-cortex distance in living cells by super-resolution microscopy. J Phys D Appl Phys 50(6):064002. https://doi.org/10.1088/1361-6463/aa52a1.

31. Crittenden JR, Bergmeier W, Zhang Y, Piffath CL, Liang Y, Wagner DD, Housman DE, Graybiel AM. (2004) CalDAG-GEFI integrates signaling for platelet aggregation and thrombus formation. Nat Med 10(9):982–6. https://doi.org/10.1038/nm1098.

32. Franke B, van Triest M, de Bruijn KM, van Willigen G, Nieuwenhuis HK, Negrier C, Akkerman JW, Bos JL. (2000) Sequential regulation of the small GTPase Rap1 in human platelets. Mol Cell Biol 20(3):779–85. https://doi.org/10.1128/MCB.20.3.779-785.2000.

33. Gutierrez-Herrero S, Fernandez-Infante C, Hernandez-Cano L, Ortiz-Rivero S, Guijas C, Martin-Granado V, Gonzalez-Porras JR, Balsinde J, Porras A, Guerrero C. (2020) C3G contributes to platelet activation and aggregation by regulating major signaling pathways. Signal Transduct Target Ther 5(1):29. https://doi.org/10.1038/s41392-020-0119-9.

34. Gutierrez-Herrero S, Maia V, Gutierrez-Berzal J, Calzada N, Sanz M, Gonzalez-Manchon C, Pericacho M, Ortiz-Rivero S, Gonzalez-Porras JR, Arechederra M, Porras A, Guerrero C. (2012) C3G transgenic mouse models with specific expression in platelets reveal a new role for C3G in platelet clotting through its GEF activity. Biochim Biophys Acta 1823(8):1366–77. https://doi.org/10.1016/j.bbamcr.2012.05.021.

35. Rehmann H, Rueppel A, Bos JL, Wittinghofer A. (2003) Communication between the regulatory and the catalytic region of the cAMP-responsive guanine nucleotide exchange factor Epac. J Biol Chem 278(26):23508–14. https://doi.org/10.1074/jbc.M301680200.

36. Jankowski W, Saleh T, Pai MT, Sriram G, Birge RB, Kalodimos CG. (2012) Domain organization differences explain Bcr-Abl’s preference for CrkL over CrkII. Nat Chem Biol 8(6):590–6. https://doi.org/10.1038/nchembio.954.

37. Maia V, Ortiz-Rivero S, Sanz M, Gutierrez-Berzal J, Alvarez-Fernandez I, Gutierrez-Herrero S, de Pereda JM, Porras A, Guerrero C. (2013) C3G forms complexes with Bcr-Abl and p38alpha MAPK at the focal adhesions in chronic myeloid leukemia cells: implication in the regulation of leukemic cell adhesion. Cell Commun Signal 11(1):9. https://doi.org/10.1186/1478-811X-11-9.

38. Vuori K, Hirai H, Aizawa S, Ruoslahti E. (1996) Introduction of p130cas signaling complex formation upon integrin-mediated cell adhesion: a role for Src family kinases. Mol Cell Biol 16(6):2606–13. https://doi.org/10.1128/MCB.16.6.2606.

39. Mitra A, Kalayarasan S, Gupta V, Radha V. (2011) TC-PTP dephosphorylates the guanine nucleotide exchange factor C3G (RapGEF1) and negatively regulates differentiation of human neuroblastoma cells. PLoS One 6(8):e23681. https://doi.org/10.1371/journal.pone.0023681.

40. Shimonaka M, Katagiri K, Nakayama T, Fujita N, Tsuruo T, Yoshie O, Kinashi T. (2003) Rap1 translates chemokine signals to integrin activation, cell polarization, and motility across vascular endothelium under flow. J Cell Biol 161(2):417–27. https://doi.org/10.1083/jcb.200301133.

41. Huang Y, Clarke F, Karimi M, Roy NH, Williamson EK, Okumura M, Mochizuki K, Chen EJ, Park TJ, Debes GF, Zhang Y, Curran T, Kambayashi T, Burkhardt JK. (2015) CRK proteins selectively regulate T cell migration into inflamed tissues. J Clin Invest 125(3):1019–32. https://doi.org/10.1172/JCI77278.

42. Park T. (2021) Crk and CrkL as Therapeutic Targets for Cancer Treatment. Cells 10(4). https://doi.org/10.3390/cells10040739.

43. Ichiba T, Kuraishi Y, Sakai O, Nagata S, Groffen J, Kurata T, Hattori S, Matsuda M. (1997) Enhancement of guanine-nucleotide exchange activity of C3G for Rap1 by the expression of Crk, CrkL, and Grb2. J Biol Chem 272(35):22215–20. https://doi.org/10.1074/jbc.272.35.22215.

44. Alonso-García N, Ingles-Prieto A, Sonnenberg A, De Pereda JM. (2009) Structure of the Calx-beta domain of the integrin beta4 subunit: insights into function and cation-independent stability. Acta Crystallogr D Biol Crystallogr 65(8):858–71. https://doi.org/10.1107/S0907444909018745.

45. Kubala MH, Kovtun O, Alexandrov K, Collins BM. (2010) Structural and thermodynamic analysis of the GFP:GFP-nanobody complex. Protein Sci 19(12):2389–401. https://doi.org/10.1002/pro.519.

46. Manso JA, Garcia Rubio I, Gomez-Hernandez M, Ortega E, Buey RM, Carballido AM, Carabias A, Alonso-Garcia N, de Pereda JM. (2016) Purification and Structural Analysis of Plectin and BPAG1e. Methods Enzymol 569:177–96. https://doi.org/10.1016/bs.mie.2015.05.002.

47. Manso JA, Gomez-Hernandez M, Carabias A, Alonso-Garcia N, Garcia-Rubio I, Kreft M, Sonnenberg A, de Pereda JM. (2019) Integrin alpha6beta4 Recognition of a Linear Motif of Bullous Pemphigoid Antigen BP230 Controls Its Recruitment to Hemidesmosomes. Structure 27(6):952–64 e6. https://doi.org/10.1016/j.str.2019.03.016.

48. Keller S, Vargas C, Zhao H, Piszczek G, Brautigam CA, Schuck P. (2012) High-precision isothermal titration calorimetry with automated peak-shape analysis. Anal Chem 84(11):5066–73. https://doi.org/10.1021/ac3007522.

49. Zhao H, Piszczek G, Schuck P. (2015) SEDPHAT--a platform for global ITC analysis and global multi-method analysis of molecular interactions. Methods 6:137–48. https://doi.org/10.1016/j.ymeth.2014.11.012.

50. Schuck P. (2000) Size-distribution analysis of macromolecules by sedimentation velocity ultracentrifugation and lamm equation modeling. Biophys J 78(3):1606–19. https://doi.org/10.1016/S0006-3495(00)76713-0.

51. Laue TM. (1996) Analytical Ultracentrifugation. Current Protocols in Protein Science 4(1):7.5.1-7.5.9. https://doi.org/https://doi.org/10.1002/0471140864.ps0705s04.

52. Dam J, Schuck P. (2005) Sedimentation velocity analysis of heterogeneous protein-protein interactions: sedimentation coefficient distributions c(s) and asymptotic boundary profiles from Gilbert-Jenkins theory. Biophysical journal 89(1):651–66. https://doi.org/10.1529/biophysj.105.059584.

53. Schuck P, Zhao H. Sedimentation Velocity Analytical Ultracentrifugation: Interacting Systems: CRC Press; 2017.

54. Svedberg T, Pedersen KO. The Ultracentrifuge: Oxford: Clarendon Press; 1940. x + 478 pp. p.

55. Beraud-Dufour S, Robineau S, Chardin P, Paris S, Chabre M, Cherfils J, Antonny B. (1998) A glutamic finger in the guanine nucleotide exchange factor ARNO displaces Mg2+ and the beta-phosphate to destabilize GDP on ARF1. EMBO J 17(13):3651–9. https://doi.org/10.1093/emboj/17.13.3651.

56. Schindelin J, Arganda-Carreras I, Frise E, Kaynig V, Longair M, Pietzsch T, Preibisch S, Rueden C, Saalfeld S, Schmid B, Tinevez JY, White DJ, Hartenstein V, Eliceiri K, Tomancak P, Cardona A. (2012) Fiji: an open-source platform for biological-image analysis. Nat Methods 9(7):676–82. https://doi.org/10.1038/nmeth.2019.

