## Supplemental figures and tables for "Crk proteins activate the Rap1 guanine nucleotide exchange factor C3G by segregated adaptor-dependent and -independent mechanisms"

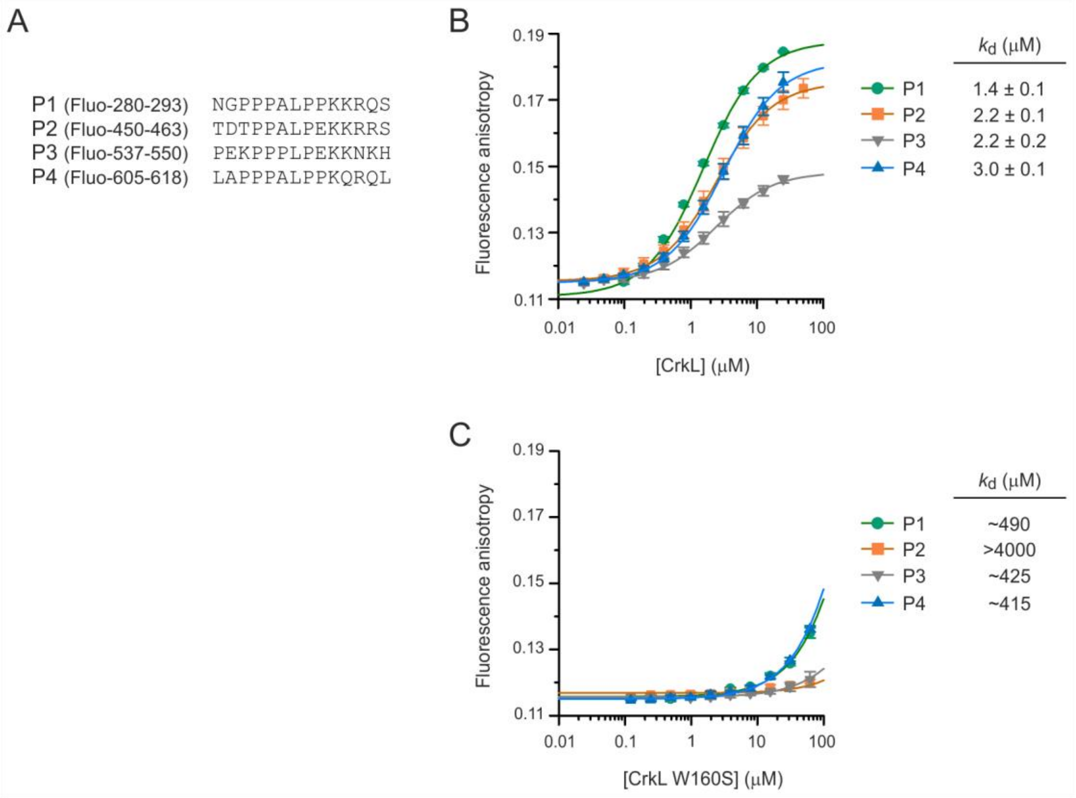


**Fig. S1. CrkL binds through the SH3N domain and with similar affinity to the four isolated PRMs of C3G**. (**A**) Sequences of four synthetic peptides that correspond to the P1 to P4 PRMs of C3G, which were used to determine the affinity of CrkL. The peptides were labeled with fluorescein at the amino-terminus. (**B**) Fluorescence anisotropy titrations of the fluorescein-labeled peptides in A (0.2 μM) with CrkL. Lines represent the 1:1 binding models fitted to the data, which yielded the apparent dissociation constants ($k_d$) ± the asymptotic standard errors. Differences in the amplitude of the anisotropy changes induced by binding of CrkL are likely to reflect variations of the microenvironment of the fluorescein probe, and are not related with the affinity. (**C**) Equivalent titrations of C3G peptides as in B but with the CrkL point mutant W160S that alters the Pro-rich binding site of the SH3N domain. Only a small fraction of the binding saturation was reached; therefore, the derived $k_d$ values are approximate only.

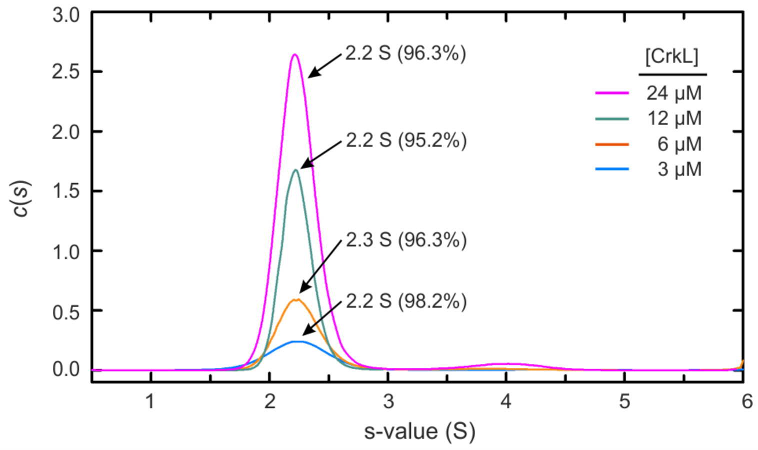


**Fig. S2. Analysis of CrkL by sedimentation velocity.** Sedimentation coefficient distributions of CrkL in 20 mM Tris-HCl, 300 mM NaCl, pH 7.5, at several loading concentrations as indicated. Over 95% of CrkL (values in parenthesis) sedimented at ~2.2 S. A minor fraction of the sample, between 1.8% and 4.8%, sedimented in a secondary *c*(s) peak at 3.7-4.3 S that is likely to correspond to CrkL dimers.

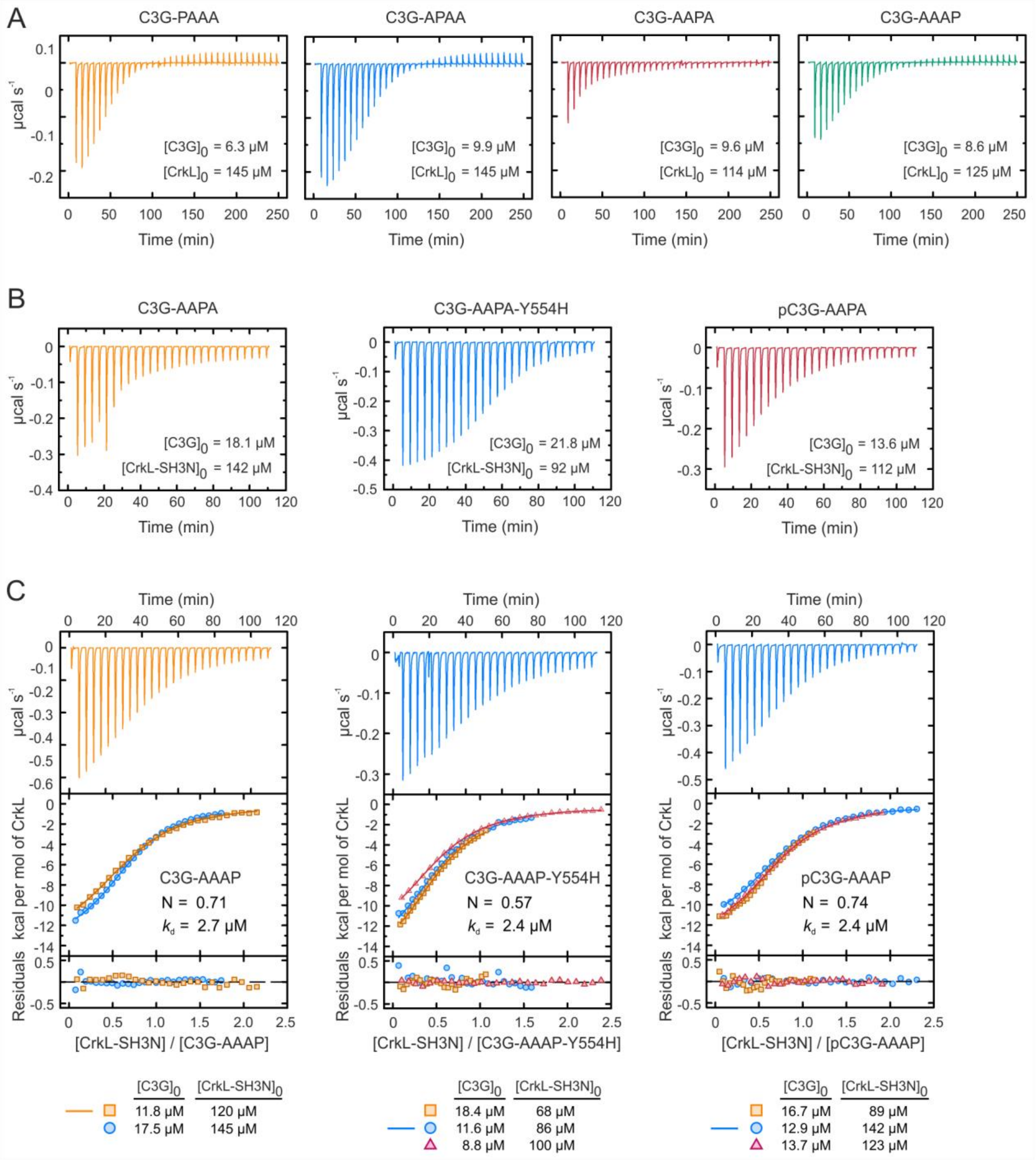


**Fig. S3. ITC analysis of the binding of CrkL to single-PRM mutants of C3G**. (**A**) Thermograms of the titration of the indicated single-PRM mutants of C3G with CrkL. The derived binding isotherms are shown in Fig. 2C. (**B**) Representative thermograms of the titrations of C3G-AAPA, the same mutant carrying the additional activating mutation Y554H, and C3G-AAPA phosphorylated with Src. The corresponding binding isotherms are shown in Fig. 2E. (**C**) ITC analysis of the binding of CrkL to C3G-AAAP, the same mutant with Y554H, and Src-phosphorylated C3G-AAAP. The upper panels correspond to representative thermograms. The values of the competent fraction of sites (N) and the microscopic $k_d$ estimated by the simultaneous global fitting of each set of three titrations are indicated.

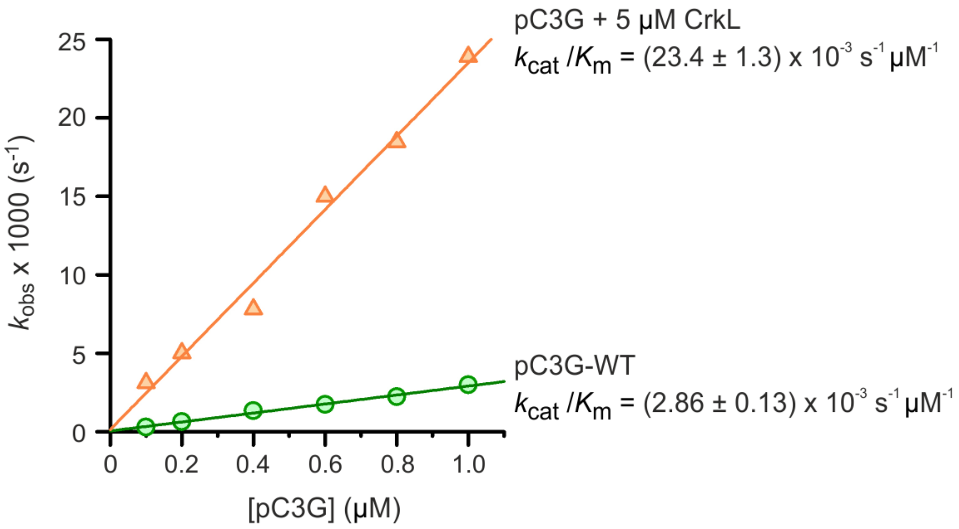


**Fig. S4. Linear concentration dependence of the nucleotide exchange activity of phospho-C3G**. Nucleotide exchange reactions were done incubating Rap1b:mant-dGDP (200 nM) with Src-phosphorylated C3G (pC3G) between 0.1 and 1 μM, alone (green circles) or in the presence of an excess of CrkL (orange triangles). The reactions were triggered by adding an excess of GDP (40 μM). The apparent dissociation rates ($k_{obs}$) were determined for each reaction by fitting a single exponential decay model. The nucleotide exchange efficiencies ($k_{cat}/K_m$) were estimated by fitting a linear dependent model (lines) to the data; and are shown as the fitted values ± the standard errors.

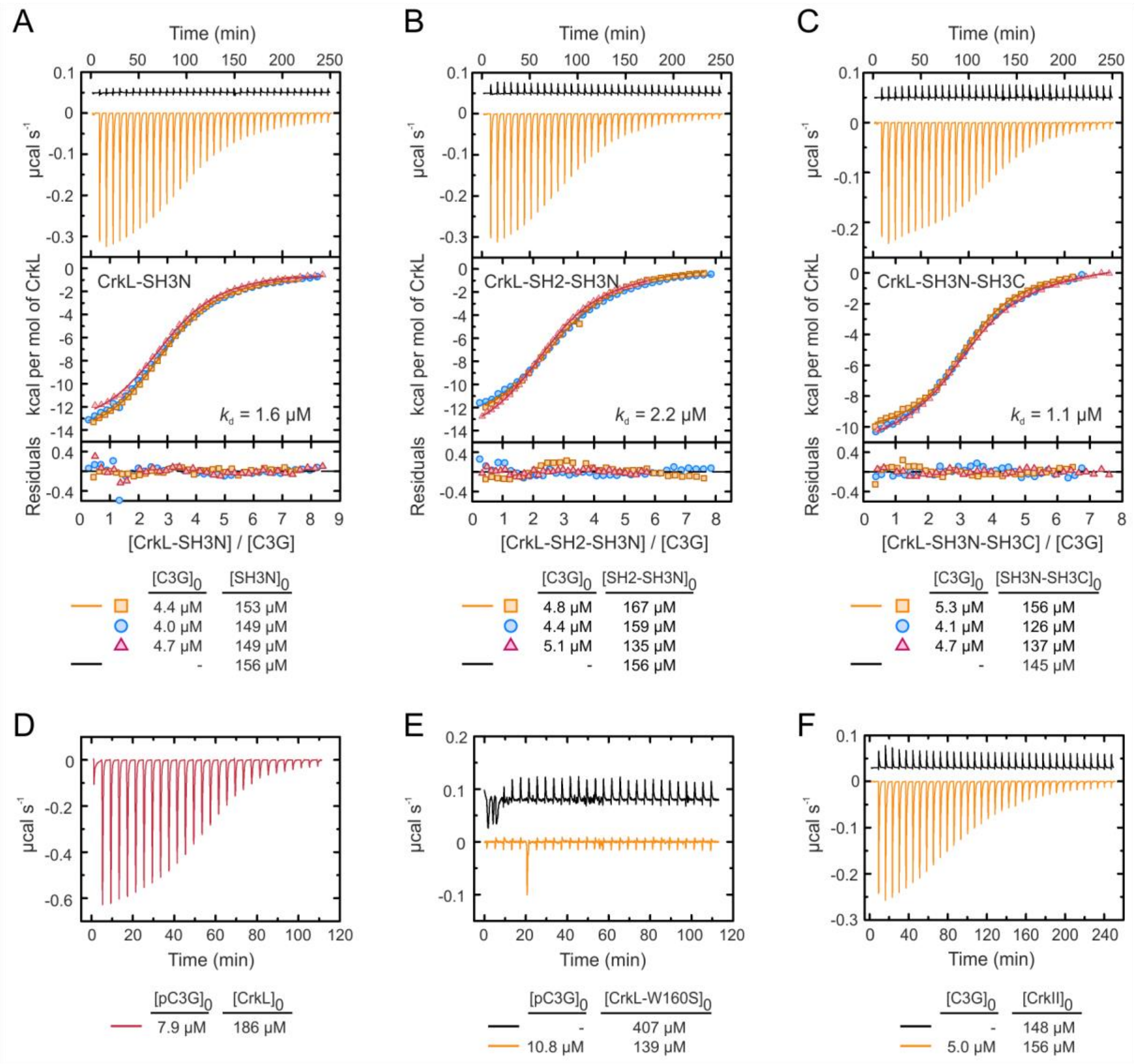


**Fig. S5. Binding of CrkL fragments and CrkII to C3G**. (**A-C**) ITC analysis of the binding of CrkL deletion mutants SH3N (A), SH2-SH3N (B), and SH3N-SH3C (C) to full-length C3G. In each figure, the upper panel shows thermograms of the dilutions of the CrkL construct in buffer (upper traces) and representative titrations of C3G with the CrkL construct (lower traces). The middle panels show the binding isotherms of three independent titrations as indicated; lines are the theoretical binding curves obtained by the global fit of model with three independent and equivalent sites. The microscopic $k_d$ values estimated by the simultaneous global fitting of each set of three titrations are indicated. Residuals from the fitted models are shown in the lower panels. The initial protein concentrations of C3G in the cell and of CrkL in the injection syringe are indicated below. (**D**) Thermogram of a representative ITC experiment of the titration of Src-phosphorylated C3G (pC3G) with CrkL. The corresponding binding isotherm is shown in Fig. 4E. (**E**) Thermograms of the dilution in buffer of CrkL-W160S (upper trace) and the titration of pC3G with CrkL-W160S (lower trace). No signal associated with binding was observed in the latter. (**F**) Thermogram of a representative ITC experiment of the titration of C3G with CrkII. The corresponding binding isotherm is shown in Fig. 4I.

**Table S1. Constructs of human C3G in the vector pETEV15b-His-Halo-TEV for expression in *E coli.***

| Construct name [a] | Mutations | N-terminal tag [b] |
|---|---|---|
| C3G | | His-Halo-TEV |
| C3G-PPAA | P540A, P541A, L543A, P544A, K546A<br>P608A, P609A, L611A, P612A, K614A | His-Halo-TEV |
| C3G-AAPP | P283A, P284A, L286, P287, P289A<br>P453A, P454A, L456A, P457A, K459A | His-Halo-TEV |
| C3G-PAAA | P453A, P454A, L456A, P457A, K459A<br>P540A, P541A, L543A, P544A, K546A<br>P608A, P609A, L611A, P612A, K614A | His-Halo-TEV |
| C3G-APAA | P283A, P284A, L286, P287, P289A<br>P540A, P541A, L543A, P544A, K546A<br>P608A, P609A, L611A, P612A, K614A | His-Halo-TEV |
| C3G-AAPA | P283A, P284A, L286, P287, P289A<br>P453A, P454A, L456A, P457A, K459A<br>P608A, P609A, L611A, P612A, K614A | His-Halo-TEV |
| C3G-AAPA-Y554H | P283A, P284A, L286, P287, P289A<br>P453A, P454A, L456A, P457A, K459A<br>Y554H<br>P608A, P609A, L611A, P612A, K614A | His-Halo-TEV |
| C3G-AAAP | P283A, P284A, L286, P287, P289A<br>P453A, P454A, L456A, P457A, K459A<br>P540A, P541A, L543A, P544A, K546A | His-Halo-TEV |
| C3G-AAAP-Y554H | P283A, P284A, L286, P287, P289A<br>P453A, P454A, L456A, P457A, K459A<br>P540A, P541A, L543A, P544A, K546A<br>Y554H | His-Halo-TEV |
| C3G-PPAP | P540A, P541A, L543A, P544A, K546A | His-Halo-TEV |
| C3G-PPPA | P608A, P609A, L611A, P612A, K614A | His-Halo-TEV |

[a] All constructs correspond to full-length C3G (residues 4-1077).
[b] The tag was removed during purification by digestion with TEV protease.

**Table S2. Oligonucleotides used to create constructs of C3G and for site directed mutagenesis.**

| Name | Sequence (5' to 3') [a] |
|---|---|
| C3Gh-004-For-NdeI | `TGAGAATTCCATATGGACTCTCAGCGTTCTCATCTCTC` |
| C3Gh-1077-Stop-Rev-BamHI | `CGGGGTACCGGATCCTAGGTCTTCTCTTCCCGG` |
| C3Gh-004-For-KKB | `TGAGGTACCGCCGCCACCATGGGATCCGACTCTCAGCGTTCTCATC` |
| C3Gh-1077-noStop-Rev-NotI | `GAATTCGCGGCCGCGGTCTTCTCTTCCCGGTC` |
| C3Gh-Y554H-For | `CAAACACATGCTGGCCCACATGCAGTTGCTGGAG` |
| C3Gh-Y554H-Rev | `CTCCAGCAACTGCATGTGGGCCAGCATGTGTTTG` |
| C3Gh-P1A-For | `GATAATGGTCCTGCAGCAGCAGCGGCACCCGCGAAAAGACAGTCGGCGC` |
| C3Gh-P1A-Rev | `GCGCCGACTGTCTTTTCGCGGGTGCCGCTGCTGCTGCAGGACCATTATC` |
| C3Gh-P2A-For | `GCAGACAGATACGGCAGCTGCTGCCGCCGAGGCGAAGCGCAGGAG` |
| C3Gh-P2A-Rev | `CTCCTGCGCTTCGCCTCGGCGGCAGCAGCTGCCGTATCTGTCTGC` |
| C3Gh-P3A-For | `GACCCAGAAAAAGCAGCTCCTGCAGCAGAGGCGAAAAACAAACACATGCTGGCC` |
| C3Gh-P3A-Rev | `GGCCAGCATGTGTTTGTTTTTCGCCTCTGCTGCAGGAGCTGCTTTTTCTGGGTC` |
| C3Gh-P4A-For | `GGCCCCGGCAGCCGCCGCAGCCCCCGCGCAGCGGCAGCTG` |
| C3Gh-P4A-Rev | `CCGCTGCGCGGGGGCTGCGGCGGCTGCCGGGGCCAGCTCC` |

[a] In the oligonucleotides used for site directed mutagenesis, the nucleotides changed are underlined in the sequence of the forward primer.

**Table S3. Constructs of human C3G in the vectors for expression in mammalian cells.**

| Construct name [a] | Vector | Mutations | C-terminal tag |
|---|---|---|---|
| C3G [b] | pLenti-C-mEGFP-IRES-BSD | | mEGFP |
| C3G-PPAA | pLenti-C-mEGFP-IRES-BSD | P540A, P541A, L543A, P544A, K546A<br>P608A, P609A, L611A, P612A, K614A | mEGFP |
| C3G-AAPP | pLenti-C-mEGFP-IRES-BSD | P283A, P284A, L286, P287, P289A<br>P453A, P454A, L456A, P457A, K459A | mEGFP |
| C3G-PAAA | pLenti-C-mEGFP-IRES-BSD | P453A, P454A, L456A, P457A, K459A<br>P540A, P541A, L543A, P544A, K546A<br>P608A, P609A, L611A, P612A, K614A | mEGFP |
| C3G-APAA | pLenti-C-mEGFP-IRES-BSD | P283A, P284A, L286, P287, P289A<br>P540A, P541A, L543A, P544A, K546A<br>P608A, P609A, L611A, P612A, K614A | mEGFP |
| C3G [b] | pEF1-mEGFP | | mEGFP |
| C3G-PPAA | pEF1-mEGFP | P540A, P541A, L543A, P544A, K546A<br>P608A, P609A, L611A, P612A, K614A | mEGFP |
| C3G-AAPP | pEF1-mEGFP | P283A, P284A, L286, P287, P289A<br>P453A, P454A, L456A, P457A, K459A | mEGFP |
| C3G-PAAA | pEF1-mEGFP | P453A, P454A, L456A, P457A, K459A<br>P540A, P541A, L543A, P544A, K546A<br>P608A, P609A, L611A, P612A, K614A | mEGFP |
| C3G-APAA | pEF1-mEGFP | P283A, P284A, L286, P287, P289A<br>P540A, P541A, L543A, P544A, K546A<br>P608A, P609A, L611A, P612A, K614A | mEGFP |

[a] All constructs correspond to full-length C3G.
[b] C3G wild type constructs were previously described [1]

**Table S4. Constructs of human CrkL and CrkII in the vectors for expression in *E coli.***

| Construct name | AA limits | Vector | Mutations | N-terminal tag |
|---|---|---|---|---|
| CrkL (full-length) | 1-303 | pETEV15b | | His-TEV [a] |
| CrkL-W160S | 1-303 | pETEV15b | W160S | His-TEV [a] |
| CrkL-SH3N | 125-182 | pETEV15b | | His-TEV [a] |
| CrkL-SH2N-SH3N | 1-182 | pETEV15b | | His-TEV [a] |
| CrkL-SH3N-SH3C | 125-303 | pETEV15b | | His-TEV [a] |
| GST-CrkL | 1-303 | pGEX-4T3-TEV | | GST-TEV |
| GST-CrkL-R39K | 1-303 | pGEX-4T3-TEV | R39K | GST-TEV |
| GST-CrkL-W160S | 1-303 | pGEX-4T3-TEV | W160S | GST-TEV |
| GST-CrkL-R39K-W160S | 1-303 | pGEX-4T3-TEV | R39K, W160S | GST-TEV |
| GST-CrkL-SH3N | 111-204 | pGEX-4T3 | | GST |
| CrkII (full-length) | 1-304 | pETEV15b | | His-TEV [a] |
| CrkL-II-L | CrkL 1-124<br>CrkII 134-191<br>CrkL 183-303 | pETEV15b | | His-TEV [a] |
| CrkII-L-II | CrkII 1-133<br>CrkL 125-182<br>CrkII 191-304 | pETEV15b | | His-TEV [a] |

[a] The poly-His tag was removed during purification by digestion with TEV protease.

**Table S5. Oligonucleotides used to create constructs of CrkL and CrkII.**

| **Name** | **Sequence (5' to 3')** [a,b] |
|---|---|
| CrkL-001-NdeI-For | TGACCATGGCATATGTCCTCCGCCAGGTTC |
| CrkL-111-SalI-For | TTTTGTCGATCTGTCTCAGCACCCA |
| CrkL-125-NdeI-For | TGACCATGGCATATGCTGGAATATGTACGGACTCTG |
| CrkL-182-Stop-BamHI-Rev | GCCGTCGACGGATCCCTACACAAGCTTTTCGACATAAGGG |
| CrkL-204-Stop-NotI-Rev | ATTAATTGCGGCCGCTCAAGCAGGTTCTGGGATCC |
| CrkL-303-Stop-BglII-Rev | GCCGTCGACAGATCTACTCGTTTTCATCTGGGTTTTGAG |
| CrkL-303-Stop-BamHI-Rev | GCCGTCGACGGATCCTACTCGTTTTCATCTGGGTTTTGAG |
| CrkL-BamHI-X-For | GGAATTCCAACAGTTATGGCATCCCAGAACCTGCTCATG |
| CrkL-BamHI-X-For | CATGAGCAGGTTCTGGGATGCCATAACTGTTGGAATTCC |
| CrkL-R39K-For | GTATGTTCCTCGTCAAGGATTCTTCCACCTGCCCTGGGG |
| CrkL-R39K-Rev | GCAGGTGGAAGAATCCTTGACGAGGAACATACCGTGGCG |
| CrkL-W160S-For | GAAGCCTGAAGAACAGTCGTGGAGTGCCCGGAAC |
| CrkL-W160S-Rev | GTTCCGGGCACTCCACGACTGTTCTTCAGGCTTC |
| CrkII-001-NdeI-For | TGACCATGGCATATGGCGGGCAACTTCGACTC |
| CrkII-304-Stop-BamHI-Rev | GCCGTCGACGGATCCTCAGCTGAAGTCCTCATCGGG |
| CrkII133-CrkL125-For | GATTCTCAGGCAGGAGGAGCTGGAATATGTACGGACTCTG |
| CrkII133-CrkL125-Rev | CAGAGTCCGTACATATTCCAGCTCCTCCTGCCTGAGAATC |
| CrkL124-CrkII134-For | CCTGCCTACAGCAGAAGATAACGCGGAGTATGTGCGAGCC |
| CrkL124-CrkII134-Rev | GGCTCGCACATACTCCGCGTTATCTTCTGCTGTAGGCAGG |
| CrkL182-CrkII192-For | GTCCCTTATGTCGAAAAGCTTGTGCCTGCCTCCGCCTCAG |
| CrkL182-CrkII192-Rev | CTGAGGCGGAGGCAGGCACAAGCTTTTCGACATAAGGGAC |
| CrkII191-CrkL183-For | TTACGTCGAGAAGTATAGAAGATCCTCACCACACGGAAAG |
| CrkII191-CrkL183-Rev | CTTTCCGTGTGGTGAGGATCTTCTATACTTCTCGACGTAA |

[a] In the oligonucleotides used for site directed mutagenesis, the nucleotides changed are underlined in the sequence of the forward primer.
[b] In the oligonucleotides used for creating the chimeric constructs, the sequences corresponding to CrkL and CrkII are shown in blue and red, respectively.

**Table S6. Oligonucleotides to subclone the abGFP4 nanobody in a modified pET22b vector.**

| **Name** | **Sequence (5' to 3')** |
|---|---|
| NbGFP-001F-Nco | GAATTCCCATGGCCCAGGTTCAACTGGTGGAAAGCGGC |
| NbGFP-116R-Xho | GCCGAATTCCTCGAGAGAGCTCACCGTCACCTGAGTCC |

**Table S7. Parameters of the CrkL/C3G interaction determined by ITC**

| Experiment | [C3G][a] (μM) | [CrkL][a] (μM) | Type of analysis | N[b] | $k_d$ (μM) [c] | ΔH (kcal/mol)[d] |
|---|---|---|---|---|---|---|
| 1 | 4.8 | 121.9 | individual [e] | 3.2 | 2.8 ± 0.1 | -14.6 |
| 2 | 5.3 | 237.2 | individual [e] | 2.8 | 3.6 ± 0.2 | -15.2 |
| 3 | 16.8 | 211.1 | individual [e] | 2.9 | 3.3 ± 0.2 | -10.5 |
| 1, 2, 3 | n.a. | n.a. | global [f] | 3 | 2.3 [1.6, 3.4] | -13.5 |

[a] Initial concentrations in the cell (C3G) and the syringe (CrkL) at the beginning of the experiment.
[b] Stoichiometry.
[c] Microscopic dissociation constant. Estimated $k_d$ are shown as the fitted value ± asymptotic standard errors (individual analyses) or as the fitted value and the limits of the asymmetric errors within a 0.95 confidence interval shown in brackets (global analysis).
[d] Binding enthalpy.
[e] Individual analyses were done with Origin ITC.
[f] Global analysis was done with SEDPHAT.

**Table S8. Hydrodynamic parameters of C3G and CrkL**

| | | | Sedimentation coefficient | | Diffusion coefficient | | |
|---|---|---|---|---|---|---|---|
| Protein | Conc. (μM) | Conc. (mg/ml) | s (S) | $s_{20,w}$ (S) | D ($m^2$ $s^{-1}$) | MW (kDa)[a] | MW/MW1[b] |
| C3G | 1.6 | 0.2 | 4.6 | 4.9 | $3.36 \times 10^{-11}$ | 127 | 1.04 |
| CrkL | 3 | 0.1 | 2.2 | 2.4 | n.d. | n.d. | n.d. |
| CrkL | 6 | 0.2 | 2.3 | 2.4 | n.d. | n.d. | n.d. |
| CrkL | 12 | 0.4 | 2.2 | 2.4 | n.d. | n.d. | n.d. |
| CrkL | 24 | 0.8 | 2.2 | 2.4 | $5.94 \times 10^{-11}$ | 34.8 | 1.03 |

[a] Mass derived from the Svedberg equation.
[b] MW1 is the theoretical mass of the monomeric proteins, derived from their sequence.

**Table S9. Parameters of the CrkL/C3G interaction determined by ITC and sedimentation velocity**

| Method | Type of analysis [a] | N[b] | $k_d$ (μM)[c] | ΔH (kcal/mol) |
|---|---|---|---|---|
| ITC | Global | 3 | 2.3 [1.6, 3.4][d] | -13.5 |
| SV | Global | 3 | 1.4 [0.7, 2.3][d] | n/a |
| ITC & SV | Global | 3 | 2.3 [1.7, 3.1][d] | -13.0 |

[a] Global analyses were done with SEDPHAT.
[b] N is the number of symmetric binding sites in the model.
[c] Microscopic dissociation constant.
[d] Numbers in brackets are the limits of the asymmetric errors within a 0.95 confidence interval.

**Table S10. Parameters of the CrkL binding to single-PRM mutants of C3G determined by ITC[a]**

| C3G | CrkL | Num exp [b] | N [c] | $k_d$ (μM) | ΔH (kcal/mol)[d] |
|---|---|---|---|---|---|
| C3G-PAAA | CrkL | 1 | 0.86 [0.84, 0.88] | 0.9 [0.8, 1.0][e] | -11.5 |
| C3G-APAA | CrkL | 1 | 0.78 [0.76, 0.80] | 1.2 [1.0, 1.4] | -13.7 |
| C3G-AAPA | CrkL | 1 | 0.13 [0.10, 0.17] | 2.1 [1.5, 3.0] | -13.5 [f] |
| C3G-AAAP | CrkL | 1 | 0.55 [0.51, 0.58] | 2.4 [2.1, 2.8] | -13.6 |
| C3G-AAPA | CrkL-SH3N | 2 | 0.20 [0.17-0.24] | 3.5 [2.7, 4.6] | -14.0 [f] |
| C3G-AAPA-Y554H | CrkL-SH3N | 3 | 0.47 [0.46-0.48] | 0.8 [0.7, 0.9] | -14.1 |
| pC3G-AAPA | CrkL-SH3N | 3 | 0.08 [0.06-0.11] | 2.1 [1.3-3.3] | -14.0 [f] |
| C3G-AAAP | CrkL-SH3N | 2 | 0.71 [0.70-0.73] | 2.7 [2.4-3.1] | -14.1 |
| C3G-AAAP-Y554H | CrkL-SH3N | 3 | 0.57 [0.55-0.58] | 2.4 [2.1-2.7] | -14.7 |
| pC3G-AAAP | CrkL-SH3N | 3 | 0.74 [0.73-0.74] | 2.4 [2.2-2.7] | -13.2 |

[a] Data were analyzed with SEDPHAT using a 1:1 heteroassociation model.
[b] Number of independent titration experiments.
[c] N is the competent fraction of C3G, which corresponds to the fraction of active binding sites or stoichiometry.
[d] Binding enthalpy.
[e] Numbers in brackets are the limits of the asymmetric errors within a 0.95 confidence interval.
[f] Values were fixed for analysis.

**Table S11. Parameters of the binding of CrkL fragments to C3G determined by ITC[a]**

| C3G | Crk protein | Num exp [b] | $k_d$ (μM) | ΔH (kcal/mol) [c] |
|---|---|---|---|---|
| C3G WT | CrkL SH3N | 3 | 1.6 [1.5-1.8][d] | -15.4 |
| C3G WT | CrkL SH2-SH3N | 3 | 2.2 [1.9, 2.5] | -14.6 |
| C3G WT | CrkL SH3N-SH3C | 3 | 1.1 [0.8, 1.4] | -11.7 |
| pC3G WT | CrkL | 3 | 1.3 [1.2, 1.4] | -11.5 |
| C3G WT | CrkII | 3 | 2.7 [2.2-3.7] | -14.9 |

[a] Data were analyzed with SEDPHAT.
[b] Number of independent titration experiments.
[c] Binding enthalpy.
[d] Numbers in brackets are the limits of the asymmetric errors within a 0.95 confidence interval.

### References


1. Carabias A, Gomez-Hernandez M, de Cima S, Rodriguez-Blazquez A, Moran-Vaquero A, Gonzalez-Saenz P, Guerrero C, de Pereda JM. (2020) Mechanisms of autoregulation of C3G, activator of the GTPase Rap1, and its catalytic deregulation in lymphomas. Sci Signal 13(647):eabb7075. https://doi.org/10.1126/scisignal.abb7075.
